# Region-specific patterns of sexual shape variation in the human bony labyrinth: 3D geometric morphometric analysis of a sample with known genomic sex

**DOI:** 10.64898/2026.08.19.745177

**Authors:** Lumila Paula Menéndez, María Clara López-Sosa, Gustavo Daniel Montiel Hernández, Wara Siles, Hans Groh, Cassandra Rios, Candela Acosta Morano, Daniela Guevara, Paula Novellino, Daniela Mansegosa, Horacio Chiavazza, Sebastian Giannotti, Sebastian Pastor, Luis Tissera, Andrea Recalde, Ivan Diaz, María Solange Grimoldi, Eva Peralta, Cinthia Abbona, Micaela Victoria Tappatá, Mariano Del Papa, Mónica Berón, Eliana Lucero, Pablo Messineo, Mariela Gonzalez, Nahuel Scheifler, Ana Solari, Sergio Monteiro Da Silva, Anne-Marie Pessis, Ramiro Barberena, Nicolas Rascovan, Pierre Luisi, Christine Chappard

## Abstract

The human bony labyrinth has attracted increasing interest because of its taxonomic, evolutionary, and functional significance. Although sexual dimorphism has been reported in several aspects of the temporal bone, the extent to which sex, age, size, and allometry contribute to labyrinth shape variation remains poorly understood. Here, we investigated patterns of sexual shape variation in the human bony labyrinth using three-dimensional geometric morphometrics in a sample of 98 archaeological individuals from South America with known genomic sex. Centroid size and allometric effects were assessed in a subset of 90 individuals with comparable metric scaling. In addition to analysing the complete labyrinth, the cochlea and semicircular canals were examined separately to evaluate region-specific patterns of sexual shape variation. Principal Component Analysis showed extensive overlap between females and males, and overall labyrinth shape did not differ significantly between sexes. Males exhibited significantly larger labyrinths than females, and centroid size explained a small but significant proportion of overall shape variation. Regional analyses showed no evidence of significant sexual shape differences in the cochlea or in any individual semicircular canal when analysed separately. In contrast, the combined semicircular canal system exhibited subtle but significant sexual shape variation independent of centroid size, whereas morphological disparity did not differ between sexes. The geometric comparison of the female and male consensus configurations further showed that sexual shape variation was regionally heterogeneous. Whereas the cochlea exhibited a pattern of localized changes with low directional coherence, the semicircular canals displayed more coordinated regional shape changes. The male consensus also exhibited slightly higher canal circularity across all three semicircular canals, particularly the posterior canal, while differences in canal-plane orientation remained minimal. These findings demonstrate that sexual shape variation in the human bony labyrinth is subtle and anatomically partitioned among its components. Although significant sex differences in centroid size were detected across most anatomical regions, overall labyrinth shape and cochlear morphology were primarily influenced by allometry, whereas significant sex-related shape differences were detected only when the semicircular canals were considered as an integrated anatomical system. These findings demonstrate that sexual dimorphism in the human bony labyrinth is subtle but regionally heterogeneous, with the cochlea and semicircular canals exhibiting distinct patterns of shape variation, suggesting that these structures are influenced by different developmental, functional, and evolutionary processes.

## INTRODUCTION

Sexual dimorphism refers to systematic phenotypic differences between males and females of the same species arising from interactions among genetic, developmental, hormonal, functional, and environmental factors (Plavcan, 2001). In humans and other primates, these differences are expressed across a broad range of traits, including body size, skeletal morphology, soft tissue distribution, vocal characteristics, and secondary sexual features (Plavcan, 2001). Skeletal sexual dimorphism has been extensively documented in both the cranial and postcranial skeleton and has provided the foundation for much of our understanding of human biological variation (Martin & Saller, 1959; Ruff, 2000; Ubelaker & DeGaglia, 2017). Most of these differences become more pronounced during puberty under the influence of sex hormones and are therefore more evident in adults than in subadults (Ruff, 2000; White & Folkens, 2005).

While human sexual dimorphism has been extensively investigated in the cranial and postcranial skeleton, the petrous portion of the temporal bone, and particularly the bony labyrinth, has received comparatively less attention (Hardy, 1938; Osipov et al., 2013; Braga et al., 2019). This structure is of particular interest because it undergoes early ossification, exhibits remarkable developmental stability, and is protected by the petrous bone, one of the densest bones in the human skeleton, resulting in exceptional preservation in archaeological and fossil contexts (Jeffery & Spoor, 2004; Lebrun et al., 2010; Ponce de León et al., 2018; Spoor et al., 2007). Consequently, it has become an important source of information for studies of phylogeny, population history, taxonomy, and human evolution (Beaudet et al., 2019; Gunz et al., 2012; Martin et al., 2026; Ponce de León et al., 2018; Smith et al., 2025; Urciuoli et al., 2025).

Far fewer studies have examined whether the bony labyrinth also exhibits sexual dimorphism. Existing evidence suggests that sex-related differences may occur in the cochlea, vestibule, and semicircular canals (Boucherie et al., 2021; Braga et al., 2019; Hardy, 1938; Marcus et al., 2013; Osipov et al., 2013; Sato et al., 1991; Uhl et al., 2020). However, reported patterns remain inconsistent. Early studies based on linear measurements reported greater cochlear dimensions in males (Hardy, 1938; Sato et al., 1991), whereas CT-based analyses identified differences in several cochlear and vestibular measurements (Marcus et al., 2013). In contrast, Osipov et al. (2013) found evidence of sexual dimorphism primarily in the semicircular canals, while Braga et al. (2019) argued that cochlear shape itself is sexually dimorphic from birth. More recent studies have suggested that the magnitude and anatomical distribution of sexual dimorphism vary among populations and that its utility for sex estimation may be more limited than initially proposed (Boucherie et al., 2021; Uhl et al., 2020). Likewise, Ward et al. (2020) found no clear evidence of sexual dimorphism, highlighting the influence of methodological differences, sample composition, and population-specific variation. The variability reported among previous studies likely reflects a combination of biological and methodological factors. These include differences in the magnitude of sexual dimorphism across populations, the method used for sex estimation (morphological, archival, or genetic), and the analytical framework employed, ranging from linear measurements to geometric morphometrics and from analyses of the entire bony labyrinth to individual anatomical components. In addition, previous studies have quantified different aspects of morphology, with some analysing overall form (i.e., the combined effects of size and shape), whereas others examined shape and size separately. These methodological differences make direct comparisons among studies difficult and may partly explain the contrasting patterns of sexual dimorphism reported in the literature.

Interpreting these patterns also requires considering the distinct developmental and functional roles of each component of the inner ear. The cochlea transduces sound into neural signals (Manoussaki et al., 2008; Spoor et al., 2007), whereas the semicircular canals detect angular acceleration and stabilize gaze through the vestibulo-ocular reflex (Angelaki & Cullen, 2008). The vestibule, comprising the utricle and saccule, detects linear acceleration and head position, contributing to balance and postural control (Curthoys et al., 2017; Goldberg, 2022). Together, these structures form an integrated sensory system, yet their distinct developmental trajectories and functional specializations suggest that morphological variation, including sexual dimorphism, may not be expressed uniformly across the components of the bony labyrinth. This interpretation is also consistent with previous studies suggesting that different regions of the bony labyrinth capture distinct aspects of morphological and evolutionary variation (Le Maître et al., 2017).

Physiological evidence further indicates that males and females differ in aspects of both auditory and vestibular function. Females generally exhibit greater sensitivity to high-frequency sounds and stronger cochlear responses, whereas males may show enhanced sensitivity at lower frequencies (Bilger et al., 1990; Guimaraes et al., 2006; McFadden, 1993). These differences have been linked to hormonal influences on cochlear physiology and auditory processing (Kim et al., 2002). Likewise, increasing evidence suggests that vestibular physiology may also exhibit sex-related variation. Several vestibular disorders show sex differences in prevalence, and hormonal influences have been proposed as one mechanism underlying these differences, although studies of vestibular-evoked myogenic potentials and vestibulo-ocular reflexes have generally reported subtle or inconsistent sex differences (Smith et al., 2019). Nevertheless, it remains uncertain whether these physiological differences are accompanied by consistent morphological variation in the osseous components of the bony labyrinth. Given the early ossification and developmental stability of the bony labyrinth (Jeffery & Spoor, 2004), any morphological expression of sexual dimorphism is expected to be subtle and potentially shaped by developmental constraints, population history, and methodological differences among studies.

To address these gaps, we investigate sexual dimorphism in the bony labyrinth using an archaeological sample of Middle to Late Holocene individuals from South America, all with associated genomic sex estimates. We assess size and shape variation in the cochlea and semicircular canals using high-resolution three-dimensional geometric morphometrics while explicitly evaluating the influence of age on observed patterns of sexual variation. This represents the first study of inner ear sexual dimorphism in South American populations combining genomic sex determination with landmark-based high-resolution geometric morphometrics. By combining these methodological approaches, this study minimizes a major source of uncertainty that has limited previous investigations of inner ear sexual dimorphism.

The present study is designed to characterize general patterns of sexual variation across a heterogeneous archaeological sample rather than to estimate population-specific patterns of sexual dimorphism. By incorporating individuals from different geographic and chronological contexts, population-related variation is retained as part of the background range of human morphological variability against which sex-related differences are evaluated. Population-specific analyses would require larger and more balanced samples within geographic and temporal groups and therefore fall beyond the scope of the present study.

Our aim is to characterize the magnitude and anatomical distribution of sexual dimorphism across the components of the bony labyrinth and to assess its biological significance within a developmentally constrained anatomical system. We hypothesize that sexual shape variation is not uniformly distributed throughout the bony labyrinth and that the cochlea and semicircular canals exhibit distinct patterns of variation owing to their different developmental and functional histories. By using chromosomal rather than morphologically estimated sex, this study provides a robust framework for testing these patterns. More broadly, this study provides a framework for investigating subtle patterns of morphological variation in one of the earliest-forming and most developmentally stable regions of the human skeleton.

## 2. MATERIALS AND METHODS

### 2.1 Archaeological samples and selection criteria

The study sample comprises Middle-to-Late Holocene human remains from archaeological sites in Argentina, Chile, and Brazil (Table 1; Table S1-S2). Individuals were selected based on two criteria:

1. preservation of at least one osseous labyrinth suitable for three-dimensional reconstruction, and
2. the availability of an independent genomic sex determination.

**Table 1.**
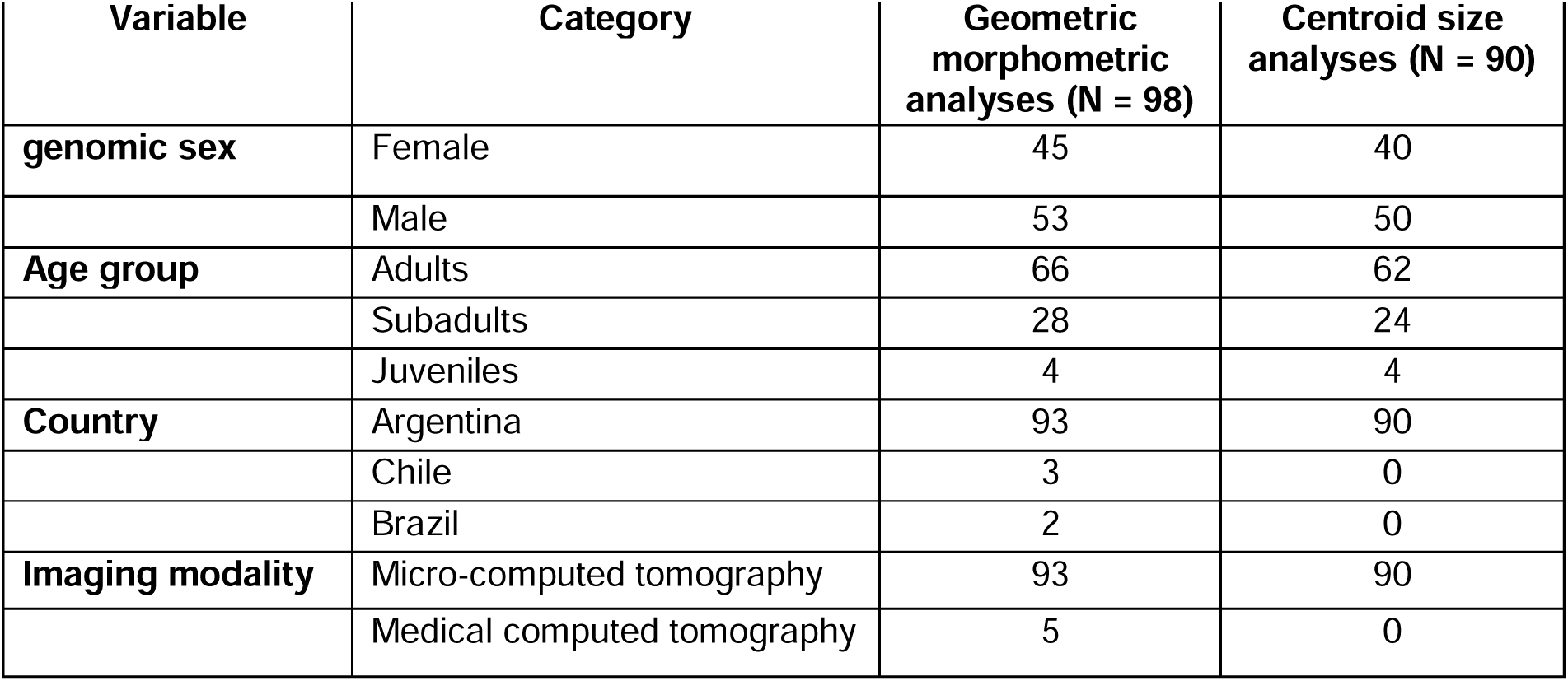
Summary of the archaeological sample included in this study. Shape analyses were performed on the complete analytical sample (N = 98). Centroid size analyses were restricted to the 90 Argentine individuals scanned using the same micro-computed tomography protocol (Table S1).

| Variable | Category | Geometric morphometric analyses (N = 98) | Centroid size analyses (N = 90) |
| --- | --- | --- | --- |
| **genomic sex** | Female | 45 | 40 |
|  | Male | 53 | 50 |
| **Age group** | Adults | 66 | 62 |
|  | Subadults | 28 | 24 |
|  | Juveniles | 4 | 4 |
| **Country** | Argentina | 93 | 90 |
|  | Chile | 3 | 0 |
|  | Brazil | 2 | 0 |
| **Imaging modality** | Micro-computed tomography | 93 | 90 |
|  | Medical computed tomography | 5 | 0 |

The final sample consisted of 98 individuals with known genomic sex, including 45 chromosomally female and 53 chromosomally male individuals. Based on the osteological age estimates available from the contributing collections, 66 individuals were classified as adults (>20 years), 28 as subadults (0–12 years), and 4 as juveniles (12–20 years) (Table 1). Age categories followed the original osteological assessments provided by each collection. Because age estimates varied among collections, with some reporting estimated age ranges and others broader categorical assignments, all analyses were performed using these standardized age groups.

Most individuals derive from archaeological sites distributed across Argentina, including the Andes, Cuyo, Central Argentina, the Paraná Delta, and the Pampas (Table S1-S2; Barberena et al., 2017, 2026; Berón, 2018; Chiavazza et al., 2015; Cigliano et al., 1973; Cocco et al., 2004; Del Papa et al., 2020; Díaz & Recalde, 2025; Durán & Novellino, 1999-2000; Durán et al., 2018; Gil et al., 2020; Guevara et al., 2022; Lehmann-Nitsche, 1910; Peralta et al., 2024; Politis & Bonomo, 2011; Ramos Van Raap & Bonomo, 2016; Recalde et al., 2024, 2026; Rivero et al., 2015; Rusconi, 1962; Scheifler et al., 2024; Spano et al., 2014; Tissera, 2014, 2024; Tissera et al., 2019). Most Argentine samples, with the exception of three from the Pampas (Table S1-S2), were previously collected as part of a paleogenomics project led by NR at the Institute Pasteur, France, and studied together by following an interdisciplinary and sustainable approach that combined imaging and genetics analyses (Barberena et al., 2026; Menéndez et al., 2026). The other three samples from the Pampas include individuals from Laguna Chica and Arroyo Seco 2, which are currently curated at the Instituto de Investigaciones Arqueológicas y Paleontológicas del Cuaternario Pampeano (INCUAPA), Olavarría, Argentina (Politis et al., 2014; Scheifler et al., 2025).

The sample also includes one individual from Última Esperanza (southern Chile), two individuals from Chilean Tierra del Fuego, and two individuals from Serra da Capivara (Piauí, northeastern Brazil) (Table S1). The Brazilian individuals, from Toca dos Caboclos and Tenente Luiz, are curated at the Fundação Museu do Homem Americano (FUMDHAM), Brazil (Cunha, 2014; Guidon et al., 1998; Mendonça de Souza et al., 2002). The Tierra del Fuego individuals are part of the Rousson et Willems collection, housed at the Musée de l’Homme, Paris, France (Galland & Friess, 2016), whereas the Última Esperanza individual is curated at the University of Zurich, Switzerland (Menéndez et al., 2025). A 3D model of the latter is publicly available for research purposes (Schmelzle et al. (2025).

All analyses were conducted under the corresponding institutional permissions and research agreements. Detailed information on the archaeological site, chronology, geographic region, estimated age, genomic sex, and imaging method for each individual is provided in Supplementary Table S1.

### 2.2 Computed tomography imaging

The osseous labyrinths were reconstructed from either micro-computed tomography (µCT) or medical computed tomography (CT) scans, depending on sample availability, logistics, and institutional facilities (Table S1). Most samples were µCT-scanned by LPM, a few were µCT-scanned in local facilities by specialized technicians (Última Esperanza, Tierra del Fuego), whereas a smaller number were acquired using medical CT scanners operated by local imaging technicians following standardized acquisition protocols defined for this project (Medical CTs from the Argentinean Pampas and Brazil).

The samples µCT-scanned by LPM, as part of a sustainable interdisciplinary workflow (Menéndez et al., 2026), were acquired using a Bruker SkyScan 1176 micro-CT scanner at the Faculty of Medicine, Université Paris Cité (formerly Paris Diderot University). Scanning parameters included an isotropic voxel size of 35 μm, a voltage of 65 kV, a current of 393 μA, an exposure time of 132 ms, and a 0.5-mm aluminium filter.

The µCT-scans from Tierra del Fuego were acquired at the AST-RX Platform, Muséum national d’Histoire naturelle (Paris, France), using a GE Phoenix v|tome|x L 450 micro-computed tomography (µCT) system. Scanning was performed with an X-ray voltage of 160 kV, an X-ray current of 800 µA, a 0.9-mm copper filter, an exposure time of 1000 ms per projection, and a total of 2600 projections, resulting in an isotropic voxel size of 128.5 µm.

The µCT-scanned sample from Última Esperanza was acquired using a Nikon XTH 225 ST high-resolution micro-computed tomography (µCT) system at the Departments of Palaeontology and Anthropology, University of Zurich (Menéndez et al., 2025; Schmelzle et al., 2025). Scanning was performed using the following acquisition parameters: voxel size of 12 μm, X-ray voltage of 180 kV, X-ray current of 298 μA, and a 1-mm-thick copper filter.

The medical CT-scans of the Laguna Chica and Arroyo Seco 2 samples from the Argentinean Pampas were acquired at local medical imaging facilities. The Laguna Chica samples were scanned at Sanatorio CEMEDA, Olavarría, Argentina, using a Neuviz 16 multidetector CT scanner (PNMS). Scanning was performed at 90 kV and 40 mA. The reconstructed images had an in-plane pixel size of 1.088 × 1.088 mm and comprised 389 axial slices. The Arroyo Seco 2 sample was scanned at the Hospital Municipal Dr. Héctor M. Cura, Olavarría, Argentina, using a Philips Ingenuity CT scanner. The acquisition parameters were 120 kV, 244 mA, a slice thickness and spacing of 0.67 mm, an in-plane pixel size of 0.260 × 0.260 mm, and a reconstruction matrix of 768 × 768 pixels.

The medical CT-scans from Brazil were scanned at Clínica SIR, Recife, Brazil, using a GE Brivo CT385 Series medical computed tomography (CT) scanner (GE Medical Systems). The acquisition parameters included a slice thickness of 0.625 mm, an in-plane pixel size of 0.488 × 0.488 mm, and a reconstruction matrix of 512 × 512 pixels.

Although image resolution varied among samples, all datasets provided sufficient anatomical detail for reliable reconstruction of the cochlea, vestibule, and semicircular canals. Samples showing extensive taphonomic damage or insufficient image quality were excluded from analyses of the affected anatomical structures.

### 2.3 Genomic sex assessment

Genomic sex was determined independently of skeletal morphology using previously generated ancient DNA data. Using genomic rather than morphologically estimated sex avoids circularity when investigating skeletal sexual dimorphism and enables the inclusion of individuals for whom morphological sex estimation is unreliable or not possible, particularly subadults and fragmentary remains.

For the samples scanned at Université Paris Cité (N=90), genomic sex was determined as part of ongoing palaeogenomic projects led by NR using standard ancient DNA approaches based on the relative representation of X and Y chromosome sequence reads relative to the autosomes. Genomic sex was inferred for genomes with a nuclear Depth of Coverage (DoC) greater than 0.0001X by comparing the DoC observed in X and Y chromosomes relative to what obtained for the autosomes, following the procedure described in the Supplementary Information (Section 5.5) of Barberena et al. (2026). Genomic sex assignments for the Brazilian samples were generated and kindly shared by Ron Pinhasi, Olivia Cheronet, and David Reich. Previously published genomic sex determinations were used for three individuals from the Argentinean Pampas (Roca-Rada et al., 2021), Tierra del Fuego (Raghavan et al., 2015), and Última Esperanza (Menéndez et al., 2025). In these cases, genomic sex determinations followed the procedures described in the original publications.

### 2.4 Three-dimensional reconstruction and landmark acquisition

CT data were imported into 3D Slicer (v. 5.8.1; Fedorov et al., 2012) using either the DICOM module (for medical CTs) or the ImageStacks module (for µCTs) from the SlicerMorph extension (Rolfe et al., 2021), depending on the format of the original scans. For high-resolution or particularly large volumes, they were reduced to the region of interest prior to segmentation. This was done either through the cropping functionality within ImageStacks or using the Crop Volume module, retaining only the anatomical area relevant for subsequent processing. The osseous labyrinth was segmented in the Segment Editor using a combination of intensity thresholding and manual editing. When both temporal bones were available, the left osseous labyrinth was preferentially segmented. If the left labyrinth was absent, incompletely preserved, or obscured by sediment, the right labyrinth was segmented instead.

Segmentation was performed simultaneously in the axial, coronal, and sagittal planes to ensure the anatomical continuity of the cochlea, vestibule, semicircular canals, common crus, and ampullary regions. Particular attention was paid to areas affected by partial-volume effects, where boundaries between the osseous labyrinth and the surrounding petrous bone could become indistinct. Following segmentation, the resulting label map was exported and converted into a surface model using the Model Maker module, ensuring that decimation was set to zero to preserve full geometric detail. All reconstructions were visually inspected for segmentation artefacts, including discontinuities, artificial bridges, and isolated components. Minor segmentation errors were corrected manually before exporting the final models as polygon meshes (.ply). In case the right labyrinth was the one segmented, the resulting model was mirrored to the left using the Mirror tool in the Surface Toolbox module to ensure that all individuals were standardized to the same anatomical orientation. Segmentation was performed independently by multiple operators (MCLS, GMH, WS, HG, and CR) following a standardized protocol.

Morphological variation was quantified using a 3D geometric morphometric protocol specifically developed for the human osseous labyrinth. Landmarks were digitized on reconstructed surface models using MorphoDig (Lebrun, 2018). Each reconstruction was manually aligned within a fixed reference grid using three homologous anatomical landmarks. This standardized orientation provided a consistent spatial reference for the placement of landmarks, minimizing operator-dependent variation and improving repeatability. The complete landmark configuration consisted of 64 landmarks describing the morphology of the cochlea, vestibule, and semicircular canals (Figure 1). A complete description of landmark definitions and anatomical landmarks is provided in Table S3. To avoid interobserver errors, all landmark digitization was performed by a single observer (LPM).

**Figure 1.**
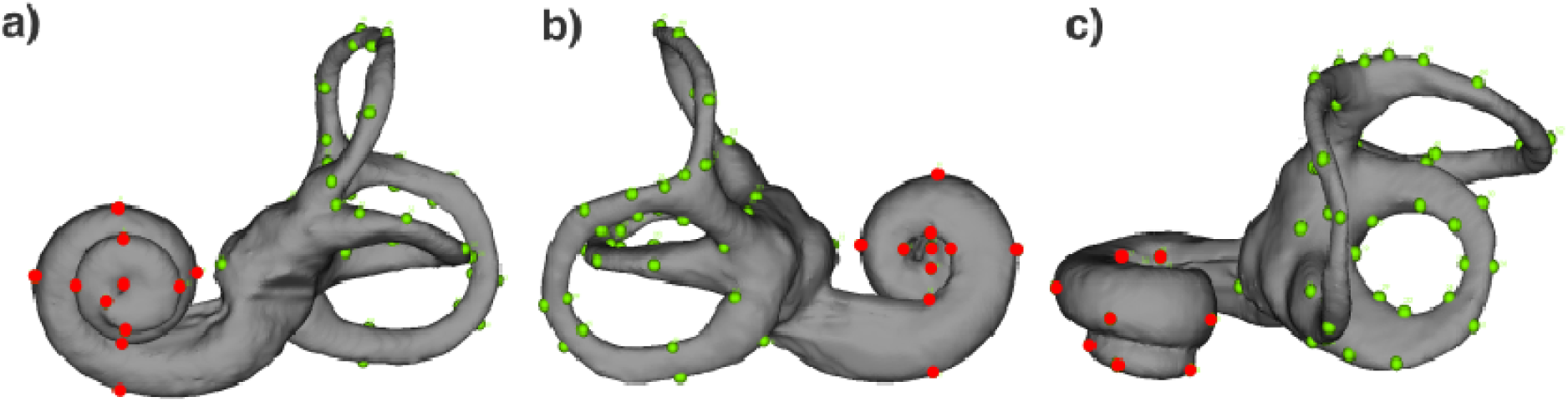
Anatomical configuration of the 64 landmarks used for the geometric morphometric analysis of the human bony labyrinth, shown in (a) anterior, (b) posterior, and (c) superior views. Detailed landmark definitions are provided in Supplementary Table S3. Landmarks belonging to the cochlea are shown in red, whereas those defining the vestibular system (vestibule and semicircular canals) are shown in green.

To investigate the anatomical distribution of sexual dimorphism, the complete landmark configuration was partitioned into biologically meaningful subsets corresponding to the cochlea and semicircular canals. While the cochlea was described using 17 landmarks, the vestibular system configuration (semicircular canals and vestibule) comprised a total of 47 landmarks (Figure 1, Table S3). From the latter, subsets were defined for the anterior, posterior, and lateral semicircular canals, allowing each canal to be analysed independently. Landmark coordinates were exported in TPS format and subsequently imported into R (version 4.4.2) for geometric morphometric and statistical analyses.

### 2.5 Geometric morphometric analyses

All geometric morphometric analyses were performed in R (version 4.4.2; R Core Team, 2025) using the packages geomorph (Baken et al., 2021), Morpho (Schlager, 2017), and associated libraries.

Landmark configurations were aligned using Generalized Procrustes Analysis (GPA) (Rohlf & Slice, 1990), which removes variation attributable to translation, rotation, and isometric scaling while preserving shape information. Consequently, shape variation reflects differences in the relative spatial configuration of landmarks that are independent of position, orientation, and overall size (Bookstein, 1991). To investigate the anatomical distribution of sexual dimorphism, independent GPAs were performed for the complete osseous labyrinth, the cochlea, the combined semicircular canals, and each semicircular canal (anterior, posterior, and lateral) analysed separately.

A small subset of samples (n = 8) acquired using different imaging workflows exhibited incompatible absolute mesh scaling, preventing direct comparison of centroid size. Because GPA removes differences in scale, these individuals were retained for all shape analyses but excluded from centroid size and allometric analyses. Consequently, analyses involving centroid size were performed on the subset of 90 individuals with comparable metric scaling.

Centroid size was calculated as the square root of the summed squared distances of all landmarks from their centroid (Bookstein, 1991). Because each anatomical configuration was analysed independently, centroid size represents the size of the corresponding anatomical structure rather than the osseous labyrinth as a whole. Since most samples consisted of isolated petrous or temporal bones rather than complete crania, an estimate of overall cranial size was not available, and centroid size was therefore used only as a measure of the size of each configuration.

### 2.6 Statistical analyses

All statistical analyses were conducted in R (R Core Team, 2025). Because different components of the bony labyrinth may be subject to distinct developmental and functional constraints, statistical analyses were performed independently for the complete bony labyrinth, the cochlea, the combined semicircular canals, and the anterior, posterior, and lateral semicircular canals. This framework allowed us to determine whether patterns of shape variation, allometry, and sexual dimorphism were expressed uniformly across the bony labyrinth or concentrated within specific anatomical components.

Morphological variation was initially explored using Principal Component Analysis (PCA) based on Procrustes shape coordinates, implemented with the gm.prcomp() function in the *geomorph* package. PCA was used solely to summarize the principal axes of shape variation and visualize morphological overlap between genetically female and genetically male individuals.

Sexual dimorphism in shape was evaluated using Procrustes Analysis of Variance (Procrustes ANOVA) implemented with the procD.lm() function in the geomorph package under the Residual Randomization Permutation Procedure (RRPP) framework. This permutation-based approach evaluates multivariate shape differences without assuming multivariate normality. Statistical significance was assessed using 10,000 random permutations. Because GPA removes isometric size variation, allometric effects were evaluated by incorporating log-transformed centroid size as a covariate in the regression models.

An initial Procrustes model including sex as the sole predictor was fitted using the complete sample (n = 98) (*Shape ∼ Sex*). To evaluate whether observed shape differences were independent of size, a second model incorporating log-transformed centroid size, sex, and their interaction was fitted using the subset of individuals with comparable metric scaling (n = 90) (*Shape ∼ log(Centroid Size) * Sex*). This model simultaneously evaluated the effects of static allometry, sex, and their interaction, allowing assessment of whether males and females differed in shape after accounting for size and whether they followed different allometric trajectories. For each model, degrees of freedom, sums of squares, coefficients of determination (R²), F-statistics, effect sizes (Z), and permutation-based P-values were recorded.

Morphological disparity was quantified as Procrustes variance using the morphol.disparity() function in the geomorph package. Differences in disparity between genetically female and male individuals were assessed using permutation tests with 10,000 iterations. In addition, regression scores derived from the allometric models were plotted against log-transformed centroid size to illustrate shape variation associated with size.

Centroid size differences between genetically female and male individuals were assessed separately for each anatomical configuration using Welch’s two-sample *t*-tests (Welch, 1947). Welch’s test was selected because centroid size analyses were restricted to individuals with comparable metric scaling, resulting in unequal sample sizes and potentially unequal variances among groups. Effect sizes for centroid size comparisons were quantified using Cohen’s *d* (Cohen, 1988). Sexual size dimorphism was expressed as the percentage by which the mean centroid size of genetically male individuals exceeded that of genetically female individuals.

To evaluate whether ontogenetic variation influenced the observed patterns of sexual shape variation, additional Procrustes ANOVAs were fitted including age group as a predictor. Models including age group alone, sex and age group jointly, and their interaction were used to assess the independent and combined contributions of age and sex to labyrinth shape.

Geographic or population affiliation was not included as a predictor in the statistical models because the archaeological sample was not designed as a balanced population-comparative dataset. Several geographic and archaeological groups are represented by small numbers of individuals, whereas others contribute substantially larger samples, precluding robust population-specific estimates of sexual dimorphism (Table S1). Accordingly, the analyses focus on sex-related variation across the pooled sample, with geographic and chronological heterogeneity treated as part of the background morphological variation.

No correction for multiple testing was applied because each anatomical configuration was analysed as an independent anatomical hypothesis.

### 2.7 Visualization and geometric characterization of sexual shape differences

To facilitate the anatomical interpretation of statistically significant shape differences, female and male consensus configurations were generated from the Procrustes-aligned landmark coordinates reconstructed using identical mesh topology. These consensus configurations were used to visualize and quantify regional patterns of sexual shape variation. Superimposed consensus models were reconstructed as dense three-dimensional surfaces to generate surface displacement heatmaps, while landmark displacement vectors were calculated directly from homologous landmarks to illustrate the magnitude and direction of local shape changes between the female and male consensus configurations. Regional geometric descriptors, including mean landmark displacement, directional coherence, angular dispersion, geometric descriptors of canal morphology (axis ratio and eccentricity), radial variation, and canal-plane orientation, were calculated to characterize differences in the spatial expression of sexual shape variation across the cochlea and semicircular canals. These analyses were descriptive and were performed to characterize the anatomical expression of shape differences identified by the inferential statistical analyses rather than to provide additional hypothesis tests.

All consensus reconstructions, geometric descriptors, and visualizations were implemented in R (R Core Team, 2025) using custom scripts based on the packages geomorph (Adams et al, 2013), Rvcg (Schlager, 2017), Morpho (Schlager, 2017), and rgl (Adler et al., 2003).

## RESULTS

### 3.1 Sexual shape variation of the whole bony labyrinth

Analyses of sexual shape variation were first performed using the full sample (n = 98) with sex as the sole predictor (Shape ∼ Sex). The PCA of the complete bony labyrinth revealed extensive overlap between genetically female and genetically male individuals in the morphospace (Figure 2). The first two principal components explained 13.5% and 10.1% of the total shape variation, respectively (23.5% cumulative variance), with no evident separation between genetically female and genetically male individuals (Figure 2).

**Figure 2.**
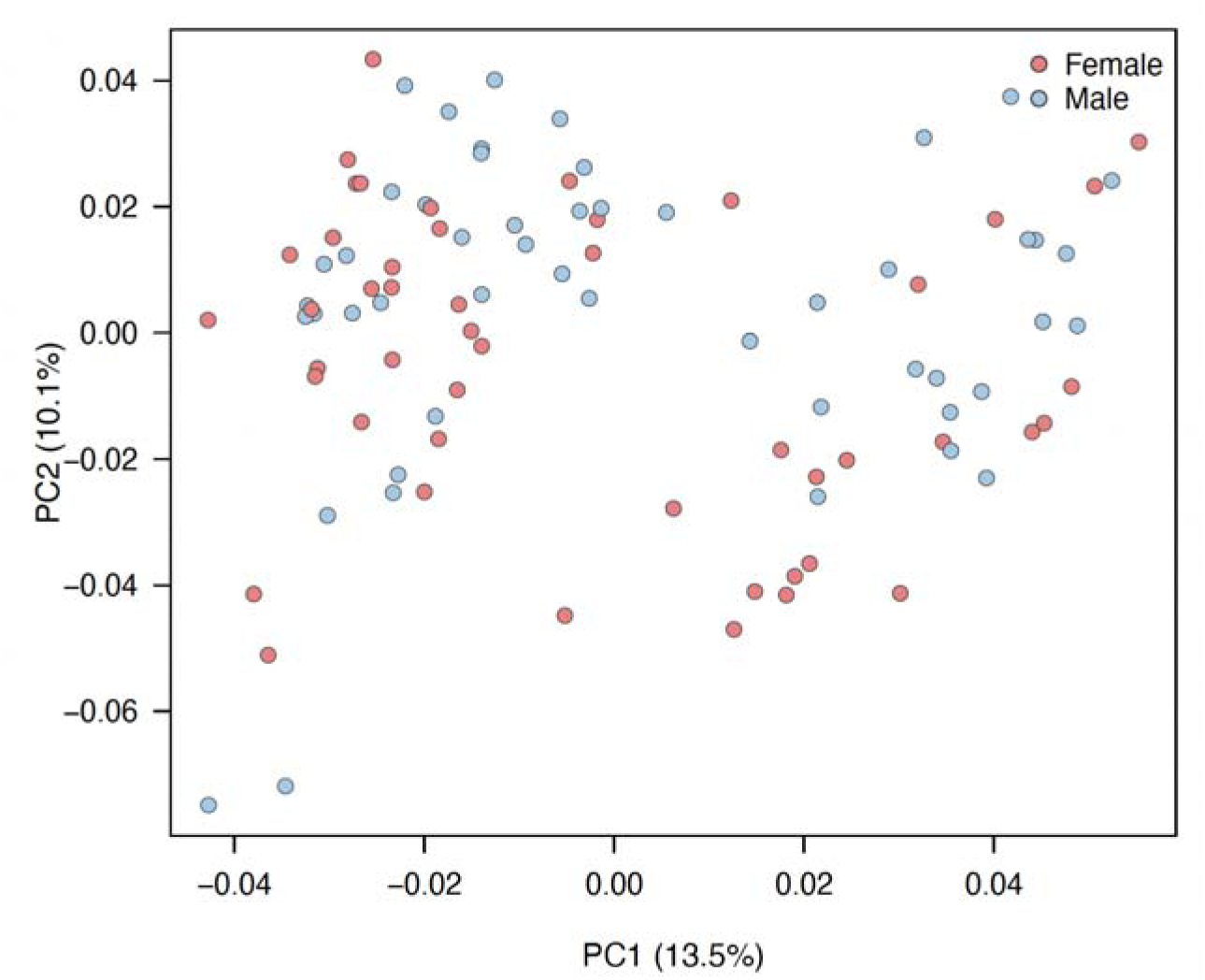
Principal component analysis of the complete bony labyrinth.

Consistent with the PCA, Procrustes ANOVA showed that sex explained only 1.48% of the total shape variation, and overall labyrinth shape did not differ significantly between genetically female and genetically male individuals (F = 1.44, R² = 0.015, p = 0.074; Table 2). Likewise, morphological disparity did not differ significantly between sexes (female Procrustes variance = 0.00573; male = 0.00598; Z = 1.47, p = 0.557), indicating comparable levels of within-sex shape variation (Table 2). Because the sample included individuals representing different ontogenetic stages, we also evaluated whether age group influenced labyrinth shape. Age group explained only 1.58% of the total shape variation and was not significantly associated with labyrinth morphology (F = 0.77, R² = 0.016, p = 0.898; Table 2). Furthermore, no significant sex-by-age group interaction was detected (F = 1.00, R² = 0.021, p = 0.443), indicating that the observed patterns of sexual shape variation were not explained by differences in age composition.

**Table 2.**
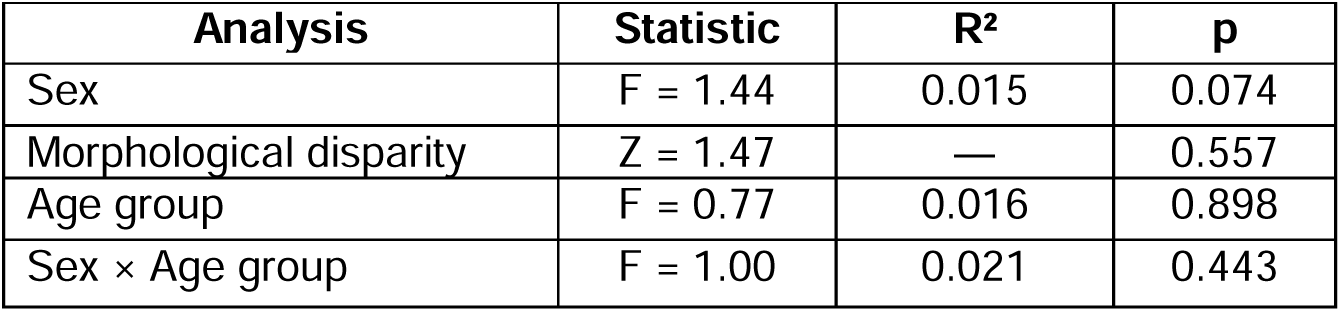
Procrustes ANOVA and morphological disparity analyses of the whole human bony labyrinth. Morphological disparity between genetically female and genetically male individuals was assessed separately using Procrustes variance, whereas shape variation was evaluated using Procrustes ANOVA.

| Analysis | Statistic | R<sup>2</sup> | p |
| --- | --- | --- | --- |
| Sex | F = 1.44 | 0.015 | 0.074 |
| Morphological disparity | Z = 1.47 | — | 0.557 |
| Age group | F = 0.77 | 0.016 | 0.898 |
| Sex × Age group | F = 1.00 | 0.021 | 0.443 |

### 3.2 Size and allometry

Because a subset of samples was acquired using a different scanning workflow that resulted in incompatible absolute mesh scaling, analyses involving centroid size were restricted to the 90 individuals with comparable metric scaling, corresponding to the individuals scanned at Paris Diderot University (Table S1). This subset was used to evaluate allometric effects and to test whether any observed sexual shape differences remained after accounting for size (Shape ∼ log(Centroid size) × Sex).

Mean centroid size was significantly larger in males (1399.5) than in females (1338.2) (Welch’s t = −5.22, p < 0.001), with a large effect size (Cohen’s d = 1.11) (Figure 3A; Table 3). Procrustes regression revealed a significant association between labyrinth shape and log-transformed centroid size, with size explaining 4.1% of the total shape variation (F = 3.74, R² = 0.041, p = 0.001). After accounting for centroid size, sex explained only 0.8% of the total shape variation and was not significant (F = 0.70, p = 0.893), indicating that overall labyrinth shape was primarily associated with size rather than sex (Figure 3B; Table 3). No significant interaction between centroid size and sex was detected (F = 0.73, R² = 0.008, p = 0.855), indicating that genetically female and genetically male individuals followed similar allometric trajectories (Figure 3B). Consistent with this result, regional analyses revealed only a weak allometric effect in the cochlea and no significant allometric effect in the semicircular canals (Supplementary Figures S3–S4).

**Figure 3.**
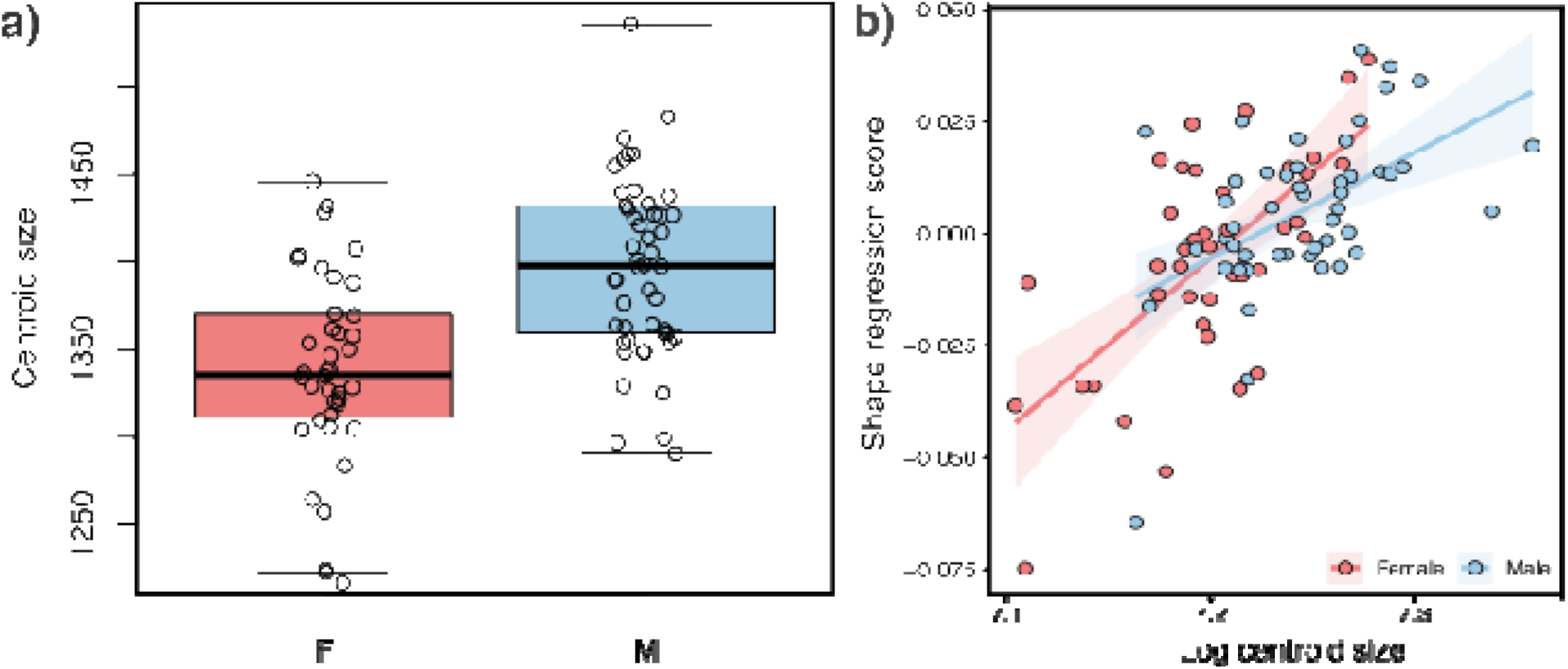
(A) Comparison of centroid size between genetically female and genetically male individuals. (B) Allometric relationship between bony labyrinth shape (represented by regression scores) and log-transformed centroid size. Lines represent linear regressions fitted separately for each sex, and shaded areas indicate 95% confidence intervals.

**Table 3.** Centroid size and allometric analyses of the whole human bony labyrinth. Differences in centroid size between genetically female and genetically male individuals were evaluated using Welch’s *t*-test, with effect size reported as Cohen’s *d*. Shape allometry was assessed using Procrustes regression (Shape ∼ log(Centroid size) × Sex), with effect size expressed as the proportion of explained shape variation (R²).

| Analysis | Statistic | Effect size | p |
| --- | --- | --- | --- |
| Centroid size | Welch's $t = -5.22$ | Cohen's $d = 1.11$ | <0.001 |
| Allometry (log centroid size) | $F = 3.74$ | $R^2 = 0.041$ | <0.001 |
| Sex (after size correction) | $F = 0.70$ | $R^2 = 0.008$ | 0.893 |
| Size × Sex interaction | $F = 0.73$ | $R^2 = 0.008$ | 0.855 |

### 3.3 Regional patterns and geometric characterization of sexual shape variation within the human bony labyrinth

To investigate whether sexual shape dimorphism was concentrated in specific anatomical regions, the cochlea and semicircular canals were analysed separately using the complete sample (n = 98) (Table 4). PCA of both regional configurations showed substantial overlap between genetically female and genetically male individuals, with no evident separation in morphospace (Supplementary Figures S1–S2).

**Table 4.** Regional analyses of sexual shape variation (Shape ∼ Sex).

| Anatomical region | F | R<sup>2</sup> | p |
| --- | --- | --- | --- |
| Cochlea | 0.68 | 0.007 | 0.807 |
| Combined semicircular canals | 1.79 | 0.018 | 0.035 |
| Anterior semicircular canal | 1.03 | 0.011 | 0.355 |
| Posterior semicircular canal | 0.90 | 0.009 | 0.483 |
| Lateral semicircular canal | 0.94 | 0.010 | 0.490 |

Consistent with the PCA, Procrustes ANOVA detected no significant sexual shape differences in the cochlea (F = 0.68, R² = 0.007, p = 0.807). In contrast, the combined semicircular canal configuration showed significant sexual shape variation, with sex explaining 1.83% of the total shape variation (F = 1.79, R² = 0.018, p = 0.035). This was the only anatomical configuration exhibiting significant sexual shape dimorphism (Table 4). Despite this difference in mean shape, morphological disparity did not differ significantly between genetically female and genetically male individuals (female Procrustes variance = 0.00962; male = 0.01049; p = 0.414), indicating comparable levels of within-sex variation.

When analysed individually, the anterior, posterior, and lateral semicircular canals showed no significant sexual shape differences (anterior: F = 1.03, R² = 0.011, p = 0.355; posterior: F = 0.90, R² = 0.009, p = 0.483; lateral: F = 0.94, R² = 0.010, p = 0.490; Table 4). The corresponding shape deformation associated with the combined semicircular canal configuration is illustrated in Figure 4.

**Figure 4.**
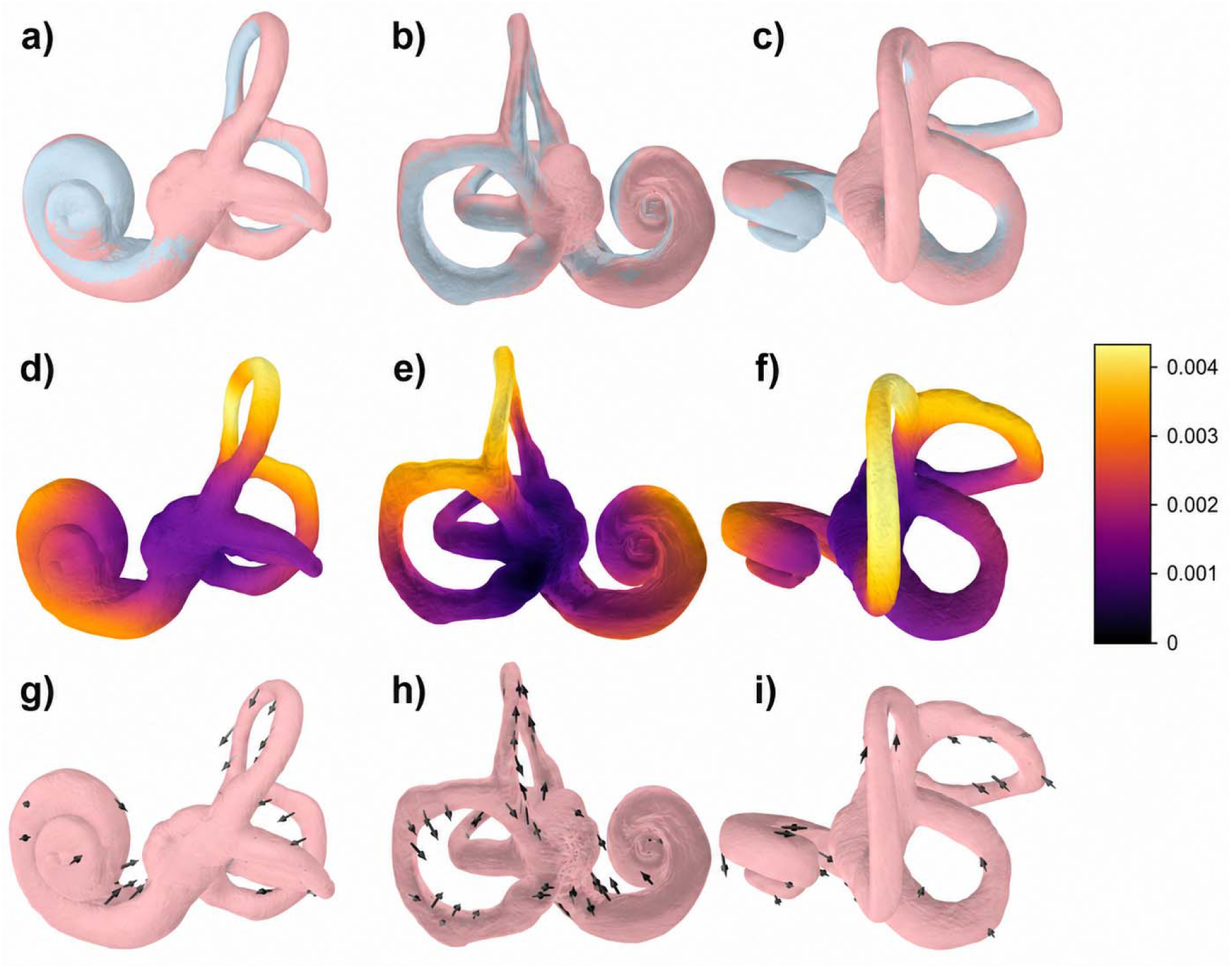
Visualization of sexual shape variation in the human bony labyrinth. Each row shows the labyrinth from the same three anatomical perspectives (anterior, posterior, and superior, from left to right). (a–c): Superimposed female (pink) and male (light blue) consensus configurations following Generalized Procrustes Analysis. (d–f): Surface displacement heatmaps illustrate the magnitude and spatial distribution of local shape differences between the female and male consensus surfaces. Colours represent vertex-wise displacement magnitude (Procrustes units), with warmer colours indicating greater displacement. (g–i): Displacement vector fields illustrating the direction and relative magnitude of local shape change from the female to the male consensus configuration. Vectors were magnified 10× for visualization.

Comparison of the female and male consensus configurations revealed that the cochlea and semicircular canals exhibit distinct patterns of sexual shape variation (Figure 4). Although the overall morphology of the bony labyrinth was highly conserved between the sexes, superimposition of the consensus models, surface displacement heatmaps, and displacement vector fields revealed subtle but regionally distributed differences in shape (Figure 4). These differences were concentrated around the junction between the cochlea and the semicircular canals and within the semicircular canals themselves, whereas the cochlea exhibited a distinct pattern of localized changes (Figure 4d-f). Visual inspection of the overlays and displacement maps further suggested that the most evident shape differences were concentrated along the superior portions of the anterior and posterior semicircular canal arches (Figure 4a-c and Figure 4g-i). Although these regional patterns were not quantified separately, they were consistent with the quantitative analyses (Table 5), which showed greater directional coherence in the semicircular canals than in the cochlea. In contrast, cochlear landmarks exhibited the lowest directional coherence and the highest angular dispersion, consistent with localized changes of the cochlear spiral rather than a uniform directional displacement. All three semicircular canals exhibited slightly higher minor-to-major axis ratios and lower eccentricity values in the male consensus, indicating a tendency toward a more circular canal morphology, with the greatest difference observed in the posterior semicircular canal (Table 5). Differences in canal-plane orientation were modest (1.1–1.6°), and the overall orientation of the labyrinth remained remarkably similar between the two consensus configurations, indicating that sexual dimorphism is primarily expressed through subtle regional modifications of canal geometry and spatial configuration rather than through major reorientation of the entire bony labyrinth.

**Table 5.** Regional patterns of shape differences between female and male consensus configurations of the human bony labyrinth. Mean landmark displacement represents the average Euclidean displacement between homologous landmarks of the female and male consensus configurations. Directional coherence quantifies the consistency of landmark displacement within each anatomical region, with higher values indicating more coordinated shape change. Mean angular deviation describes the dispersion of displacement vectors around the regional mean direction. Plane difference indicates the angular difference between the best-fitting planes of the female and male consensus configurations for each semicircular canal. Directional coherence ranges from 0 to 1, with higher values indicating more coordinated landmark displacement. Plane difference was calculated only for the semicircular canals; it is not applicable to the cochlea because it is not a planar structure.

| Region | Mean landmark displacement | Directional coherence | Mean angular deviation (°) | Plane difference (°) |
| --- | --- | --- | --- | --- |
| Cochlea | 0.00150 | 0.252 | 77.75 | – |
| Lateral semicircular canal | 0.00276 | 0.652 | 46.66 | 1.58 |
| Anterior semicircular canal | 0.00185 | 0.457 | 59.57 | 1.06 |
| Posterior semicircular canal | 0.00248 | 0.624 | 53.38 | 1.15 |

### 3.4 Regional size and allometric variation

Regional allometric analyses were performed on the subset of 90 individuals with comparable metric scaling to evaluate whether regional shape variation was associated with centroid size and whether allometric trajectories differed between genetically female and genetically male individuals.

The cochlea showed a weak but significant association between shape and log-transformed centroid size (F = 1.79, R² = 0.018, p = 0.048; Figure S3), whereas no significant allometric relationship was detected for the combined semicircular canal configuration or for the individual anterior, posterior, and lateral semicircular canals (all p > 0.20; Table 6; Figure S4). No significant interaction between log-transformed centroid size and sex was detected for the cochlea, combined semicircular canals, anterior semicircular canal, or posterior semicircular canal (Table 6). In contrast, the lateral semicircular canal exhibited a significant size-by-sex interaction (F = 1.83, R² = 0.019, p = 0.038), indicating sex-specific allometric trajectories in this structure (Table 6).

**Table 6.**
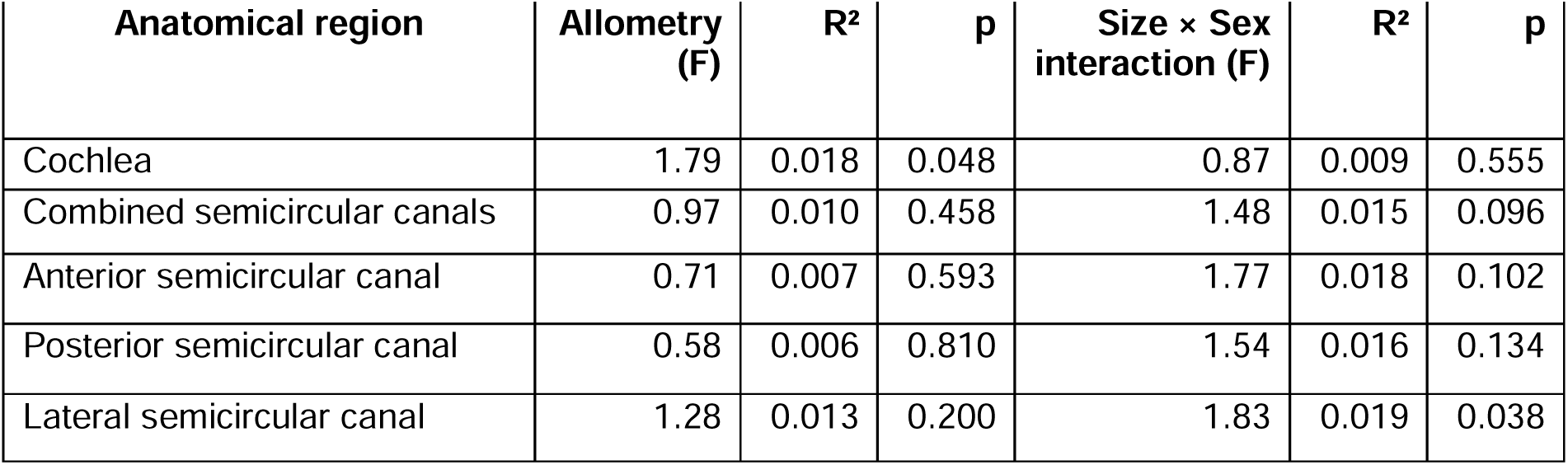
Regional allometric analyses (Shape ∼ log(Centroid Size) x Sex).

| Anatomical region | Allometry (F) | R<sup>2</sup> | p | Size × Sex interaction (F) | R<sup>2</sup> | p |
| --- | --- | --- | --- | --- | --- | --- |
| Cochlea | 1.79 | 0.018 | 0.048 | 0.87 | 0.009 | 0.555 |
| Combined semicircular canals | 0.97 | 0.010 | 0.458 | 1.48 | 0.015 | 0.096 |
| Anterior semicircular canal | 0.71 | 0.007 | 0.593 | 1.77 | 0.018 | 0.102 |
| Posterior semicircular canal | 0.58 | 0.006 | 0.810 | 1.54 | 0.016 | 0.134 |
| Lateral semicircular canal | 1.28 | 0.013 | 0.200 | 1.83 | 0.019 | 0.038 |

Centroid size analyses of the same subset revealed significant sexual size dimorphism for the whole labyrinth, cochlea, combined semicircular canals, and the anterior and posterior semicircular canals, with genetically male individuals exhibiting centroid sizes 2.6–6.3% larger than those of genetically female individuals (Table 7). No significant difference in centroid size was detected for the lateral semicircular canal (p = 0.293).

**Table 7.**
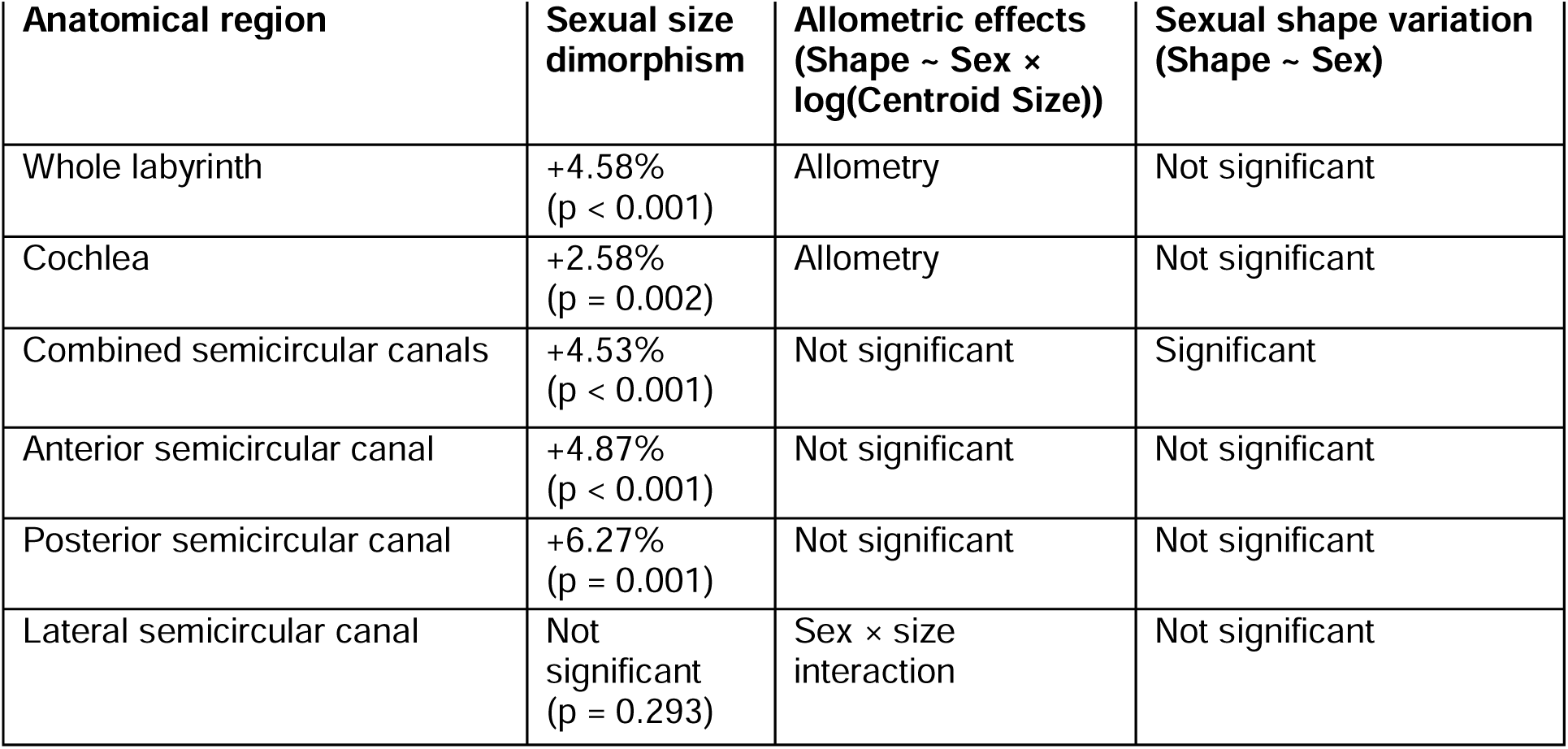
Summary of sexual size dimorphism, allometric effects, and sexual shape variation across the anatomical components of the human bony labyrinth. Centroid size differences are expressed as the percentage by which male centroid size exceeds female centroid size. *Allometric effects* summarize the results of the model *Shape ∼ Sex × log(Centroid Size)*. “Allometry” indicates a significant effect of log-transformed centroid size, whereas “Sex × size interaction” indicates significant sex-specific allometric trajectories.

Overall, the analyses reveal distinct patterns of size- and shape-related variation across the studied anatomical components of the bony labyrinth. Significant centroid size differences between genetically female and genetically male individuals were detected for most anatomical regions, whereas shape variation showed a more localized pattern. Overall labyrinth shape and cochlear morphology were primarily associated with allometric variation, whereas significant sex-related shape differences were detected only in the integrated semicircular canal system. Although no significant regional allometry was detected within the semicircular canals, the lateral semicircular canal exhibited a significant size-by-sex interaction, indicating sex-specific allometric trajectories. A summary of these findings is presented in Table 7.

## DISCUSSION

The principal finding of this study is that sexual shape variation in the human bony labyrinth is anatomically partitioned rather than uniformly distributed across the labyrinth. Although genetically male individuals exhibited significantly larger labyrinths than genetically female individuals, biological sex explained only a small proportion of overall shape variation after Procrustes superimposition. Instead, labyrinth shape was influenced primarily by allometry, and significant sexual shape differences were detected only when the semicircular canals were analysed as an integrated anatomical module. These findings demonstrate that different anatomical components of the bony labyrinth respond differently to biological sources of morphological variation, revealing a pattern of subtle, regionally heterogeneous sexual dimorphism rather than a single, global dimorphic signal.

This pattern is consistent with previous observations that the bony labyrinth is a highly conserved anatomical system (Gunz et al., 2012; Jeffery & Spoor, 2004; Spoor et al., 1994). However, our results further demonstrate that this overall conservation does not preclude subtle, anatomically partitioned sexual shape variation. Rather than being uniformly distributed across the labyrinth, sexual dimorphism was expressed through regional modifications, particularly within the integrated semicircular canal system.

Our results are consistent with those of Osipov et al. (2013), who reported that sexual dimorphism is expressed more strongly in the semicircular canals than in the cochlea. However, our findings further demonstrate that this signal is detectable only when the semicircular canals are analysed as an integrated anatomical module, whereas analyses of the individual canals revealed no significant sexual shape differences. Likewise, our findings agree with those of Boucherie et al. (2021) and Ward et al. (2020), indicating that sexual dimorphism in the human bony labyrinth is generally subtle and limited in magnitude. Although Uhl et al. (2020) suggested that the expression of sexual dimorphism may vary among populations, the present study addresses a different question: whether sex-related morphological variation can be detected across a geographically and chronologically heterogeneous archaeological sample. The detection of subtle sexual shape variation within such a heterogeneous dataset suggests that the observed signal is not restricted to a narrowly defined archaeological population. However, our sampling design does not allow us to determine whether the magnitude or anatomical expression of sexual dimorphism differs among populations. Addressing this question will require population-specific datasets with sufficiently large and balanced female and male samples.

Our results partly agree with earlier studies reporting larger cochlear dimensions in males (Hardy, 1938; Sato et al., 1991) and sex differences in cochlear and vestibular measurements (Marcus et al., 2013), as we also found significant sexual differences in centroid size across most anatomical regions. However, after separating size from shape, no significant sexual shape differences were detected in the cochlea or vestibule. It is worth noting, however, that sex differences have also been reported in longitudinal dimensions of the cochlea, including the length of the organ of Corti (Hardy, 1938) and cochlear duct length (Baguant et al., 2022), with males generally exhibiting greater lengths than females. These dimensions were not explicitly quantified in the present study and may represent additional aspects of cochlear sexual variation worth considering in future studies. Likewise, our findings differ from those of Braga et al. (2019), who reported high sex-classification accuracy based on cochlear shape alone. In contrast, our geometric morphometric analyses revealed no significant cochlear shape dimorphism after Procrustes superimposition, suggesting that the expression of sexual dimorphism depends on the morphological variables analysed and the analytical framework employed.

These contrasting findings likely reflect important methodological differences among studies. Previous investigations quantified different aspects of labyrinth morphology, with some focusing on linear dimensions (Hardy, 1938; Marcus et al., 2013; Sato et al., 1991), others analysing overall form (Osipov et al., 2013), and others restricting their analyses to individual anatomical structures, such as the cochlea (Braga et al., 2019) or the semicircular canals (Uhl et al., 2020). In contrast, the present study explicitly evaluated centroid size, shape, and allometry separately across the whole bony labyrinth and its anatomical components, allowing us to distinguish size-related from shape-related sources of variation. Furthermore, most previous studies relied on morphological or archival sex estimation (Boucherie et al., 2021; Braga et al., 2019; Osipov et al., 2013), whereas our analyses were based on independently determined genomic sex, eliminating a potential source of circularity in studies of skeletal sexual dimorphism.

An important contribution of this study comes from the consensus-based geometric characterization of sexual shape variation, which provided anatomical insight beyond that obtained from inferential statistical analyses alone. Although overall differences between the sexes were subtle, geometric characterization demonstrated that shape variation was regionally heterogeneous rather than uniformly distributed throughout the labyrinth. The cochlea exhibited a pattern of localized changes with relatively low directional coherence, whereas the semicircular canals displayed more coordinated regional shape changes. Moreover, all three semicircular canals showed a slight tendency toward increased circularity in the male consensus configuration, particularly the posterior semicircular canal, while differences in canal orientation remained minimal. Collectively, these observations indicate that sexual dimorphism is expressed primarily through localized modifications of canal geometry and spatial configuration rather than through large-scale reorganization of labyrinth morphology. These contrasting geometric patterns suggest that different labyrinth components respond differently to biological sources of morphological variation, even when the overall magnitude of sexual dimorphism remains small. Such regional differences are consistent with broader concepts of morphological integration and developmental modularity, whereby different parts of a complex anatomical structure may vary in the degree to which they respond to developmental, functional, or evolutionary influences (Klingenberg, 2008, 2014). They are also consistent with previous studies showing that the morphology of the bony labyrinth is influenced by its spatial integration within the cranial base (Le Maître, 2019). Within the bony labyrinth, one possible explanation is the developmental and functional integration of the vestibular apparatus, whose components arise through coordinated morphogenetic processes during early prenatal development and function collectively to detect angular head movements and maintain gaze and postural stability (Angelaki & Cullen, 2008; Curthoys et al., 2017; Jeffery & Spoor, 2004). Indeed, morphological distinctions for the primate bony labyrinth have been described, suggesting that a tripartite framework, comprising distinct cochlear, canalicular, and otolithic systems better reflects the structural, functional, and evolutionary complexity of the primate inner ear (Smith & Laitman, 2026). Consequently, analyses of the integrated semicircular canal system may capture coordinated patterns of variation that remain undetectable when individual canals are examined separately.

The allometric analyses provide important insight into the sources of morphological variation within the human bony labyrinth. Although genetically male individuals exhibited significantly larger labyrinths than genetically female individuals, much of the associated shape variation at the level of the whole labyrinth was explained by centroid size rather than by sex itself, highlighting the importance of accounting for allometric effects when analysing shape variation in geometric morphometrics (Klingenberg, 2016; Mitteroecker et al., 2013). After accounting for allometry, sex no longer explained a significant proportion of overall labyrinth shape variation, and no significant size-by-sex interaction was detected, indicating a common pattern of size-related shape change for the labyrinth as a whole. At the regional level, shape allometry was evident only for the cochlea, whereas the lateral semicircular canal exhibited the only significant sex-specific allometric trajectory. These findings extend previous reports of larger cochlear dimensions in males (Hardy, 1938; Marcus et al., 2013; Sato et al., 1991) by showing that sexual differences are expressed predominantly through size rather than shape and that different anatomical components of the labyrinth are influenced by distinct sources of morphological variation, consistent with broader concepts of morphological integration and modularity (Klingenberg, 2008, 2014).

The biological mechanisms underlying these subtle patterns of variation remain uncertain and are likely to involve multiple developmental and physiological processes rather than sex alone. Hormonal influences on the auditory system have been extensively documented, with estrogen and progesterone affecting cochlear mechanics, auditory thresholds, and neural processing of sound (Al-Mana et al., 2008; Hultcrantz et al., 2006; McFadden, 1993). Likewise, sex differences in hearing sensitivity and susceptibility to auditory disorders have been linked to endocrine influences throughout life, including during the menstrual cycle, pregnancy, and menopause (Guimaraes et al., 2006; Hultcrantz et al., 2006; Møller, 2012; Motlagh Zadeh et al., 2019). Although these studies concern auditory physiology rather than skeletal morphology, they suggest that sex-related biological factors may influence the development and maintenance of the inner ear. However, our results do not allow disentangling the relative contributions of genetic, hormonal, developmental, or functional factors. Determining why size-related variation predominates in the whole labyrinth and cochlea, whereas subtle sex-related differences emerge only in the integrated semicircular canal system, will require future studies integrating morphometric, developmental, physiological, and biomechanical approaches.

Beyond the study of sexual dimorphism, our findings have broader implications for interpreting morphological variation in the human bony labyrinth. The increasing use of labyrinth morphology in studies of human evolution, population history, and taxonomy (Beaudet et al., 2019; Gunz et al., 2012; Martin et al., 2026; Ponce de León et al., 2018; Smith et al., 2025; Spoor et al., 1994) has largely relied on the assumption that this structure is relatively stable and minimally influenced by sex (Jeffrey & Spoor, 2004; Spoor et al., 2007). Our results support this assumption by showing that, despite widespread sexual differences in centroid size, overall labyrinth shape is influenced more strongly by allometry than by sex, while sex-related shape variation remains anatomically localized rather than pervasive. These findings suggest that different components of the labyrinth should not necessarily be expected to respond similarly to biological or evolutionary processes. This interpretation is consistent with previous studies showing that different regions of the vestibular apparatus preserve distinct patterns of evolutionary variation (Urciuoli et al., 2020). Future studies investigating population differences, fossil hominins, or interspecific variation may therefore benefit from considering the labyrinth as a set of anatomically distinct but coordinated components rather than as a single homogeneous structure (Klingenberg, 2014; Smith & Laitman, 2026).

From an applied perspective, the limited magnitude of sexual dimorphism observed here suggests that the bony labyrinth should be used cautiously for sex estimation in archaeological and forensic contexts. Although statistically significant differences were detected, the substantial overlap between females and males indicates that labyrinth morphology alone is unlikely to provide reliable individual-level sex classification. Rather than supporting its use as a sex diagnostic, our findings highlight the value of the bony labyrinth as a model for investigating how different biological factors contribute to morphological variation across anatomically distinct components.

Several limitations should be considered when interpreting these findings. First, our sample intentionally encompasses individuals from multiple geographic and chronological contexts because the primary objective was to characterize sexual variation across a broad range of human morphological variability rather than to estimate population-specific patterns of dimorphism. However, the uneven representation of archaeological groups prevents formal evaluation of population-by-sex interactions. Consequently, the present results should be interpreted as patterns of sexual variation across the pooled sample and not as evidence that the magnitude or expression of sexual dimorphism is identical among South American populations. Population-specific variation represents an important question for future studies based on larger and more balanced comparative samples. Second, image acquisition combined micro-computed tomography and medical computed tomography. Although both modalities have been shown to produce reliable anatomical reconstructions for geometric morphometric analyses, differences in image resolution may introduce minor variation in landmark placement. Finally, although genomic sex determination is a major strength of this study because it avoids the circularity associated with morphology-based sex estimation, genomic sex represents only one aspect of biological sex and does not directly reflect the hormonal or developmental processes that contribute to skeletal morphology.

This study demonstrates that sexual shape variation in the human bony labyrinth is subtle and anatomically partitioned rather than uniformly distributed across the labyrinth. By combining genomic sex determination with three-dimensional geometric morphometrics and separate analyses of size, shape, and allometry, we provide a more comprehensive assessment of sexual dimorphism than has previously been available. Rather than representing a single homogeneous anatomical structure, the human bony labyrinth appears to comprise coordinated anatomical components that differ in their responses to biological sources of morphological variation. Our findings further suggest that these components are shaped by distinct developmental, functional, and evolutionary influences, providing a more nuanced framework for interpreting labyrinth morphology in anatomical, evolutionary, comparative, and forensic studies.

## ACKNOWLEDGEMENTS

This project was supported by two grants awarded to LPM by the Wenner-Gren Foundation: a Post-PhD Research Grant, *Human Endocranial Variation in the Southern Cone: Implications for the Peopling of South America* (Grant No. 9708), and a Hunt Postdoctoral Fellowship, *Prick Your Ears: The Contribution of Inner Ear Variation to the Peopling of South America Debate* (Grant No. 10755), as well as a grant from the German Research Foundation (DFG), *Human Morphological Diversification in the Argentinean Pampas: Implications for the Peopling of South America* (Grant No. 415489479). Those projects allowed conducting the CT-scanning of the samples included in this study. Genomic sex data was generated through the following grants awarded to NR: European Research Council ERC-2020-STG “PaleoMetAmerica” (Grant No 948800), Institut Pasteur and Centre for Scientific Research (CNRS), Unité Mixte de Recherche (UMR) 2000 funding, INCEPTION program (Investissement d’Avenir Grant ANR-16-CONV-0005). The Fundação Museu do Homem Americano (FUMDHAM) provided financial support to LPM for the dissemination of preliminary results of this study at scientific conferences.

We are grateful to Malena Vázquez and Julio Avalos from the Registro Nacional de Yacimientos, Colecciones y Objetos Arqueológicos (RENYCOA), Argentina, for their assistance with the export of the Argentine samples analysed in Paris. We thank Olivia Cheronet, Ron Pinhasi, and David Reich for generously sharing unpublished genomic sex data for the Brazilian samples included in this study. We are also grateful to Gustavo Politis, Clara Scabuzzo, Adolfo Gil, and Gustavo Neme for granting permission to study samples from the Pampas and southern Mendoza. We thank Roberto Peretti for his assistance with sample selection from the Pampas. We are grateful to Martin Friess, Véronique Laborde, and Marta Bellato for their support during the study and CT scanning of the Tierra del Fuego samples, and to Marcelo R. Sánchez-Villagra, Gabriel Aguirre Fernández, Jorge D. Carrillo Briceño, and Thomas Schmelzle for coordinating the CT scanning and segmentation of the Última Esperanza sample. Finally, we thank the imaging technicians at Hospital Municipal Dr. Héctor M. Cura, Sanatorio CEMEDA, Clínica SIR, the University of Zurich, and the AST-RX Platform for their help and assistance during CT scanning.

## AUTHORS CONTRIBUTIONS

Lumila Paula Menéndez: Conceptualization, Methodology, Data curation, Formal analysis, Investigation, Validation, Visualization, Writing-original draft, Funding acquisition, Project administration.

María Clara López-Sosa: Methodology, Data curation, Investigation, Writing-review & editing.

Gustavo Montiel Hernández: Data curation, Writing-review & editing.

Wara Siles: Data curation, Writing-review & editing.

Hans Groh: Data curation, Writing-review & editing.

Cassandra Rios: Data curation, Writing-review & editing.

Candela Acosta Morano: Data curation, Writing-review & editing.

Daniela Guevara: Resources, Writing-review & editing.

Paula Novellino: Resources, Writing-review & editing.

Daniela Mansegosa: Resources, Writing-review & editing.

Horacio Chiavazza: Resources, Writing-review & editing.

Sebastian Giannotti: Resources, Writing-review & editing.

Sebastian Pastor: Resources, Writing-review & editing.

Luis Tissera: Resources, Writing-review & editing.

Andrea Recalde: Resources, Writing-review & editing.

Ivan Diaz: Resources, Writing-review & editing.

María Solange Grimoldi: Resources, Writing-review & editing.

Eva Peralta: Resources, Writing-review & editing.

Cinthia Abbona: Resources, Writing-review & editing.

Micaela Tappatá: Resources, Writing-review & editing.

Mariano Del Papa: Resources, Writing-review & editing.

Mónica Berón: Resources, Writing-review & editing.

Eliana Lucero: Resources, Writing-review & editing.

Pablo Messineo: Resources, Writing-review & editing.

Mariela Gonzalez: Resources, Writing-review & editing.

Nahuel Scheifler: Resources, Writing-review & editing.

Ana Solari: Resources, Writing-review & editing.

Sergio Monteiro Da Silva: Resources, Writing-review & editing.

Anne-Marie Pessis: Resources, Writing-review & editing, Funding acquisition.

Ramiro Barberena: Resources, Writing-review & editing.

Nicolas Rascovan: Resources, Writing-review & editing, Funding acquisition.

Pierre Luisi: Formal analysis, Writing-review & editing.

Christine Chappard: Resources, Data curation, Writing-review & editing.

## Notes

### Competing Interest Statement

The authors have declared no competing interest.

