## Supplementary Material for "Region-specific patterns of sexual shape variation in the human bony labyrinth: 3D geometric morphometric analysis of a sample with known genomic sex"

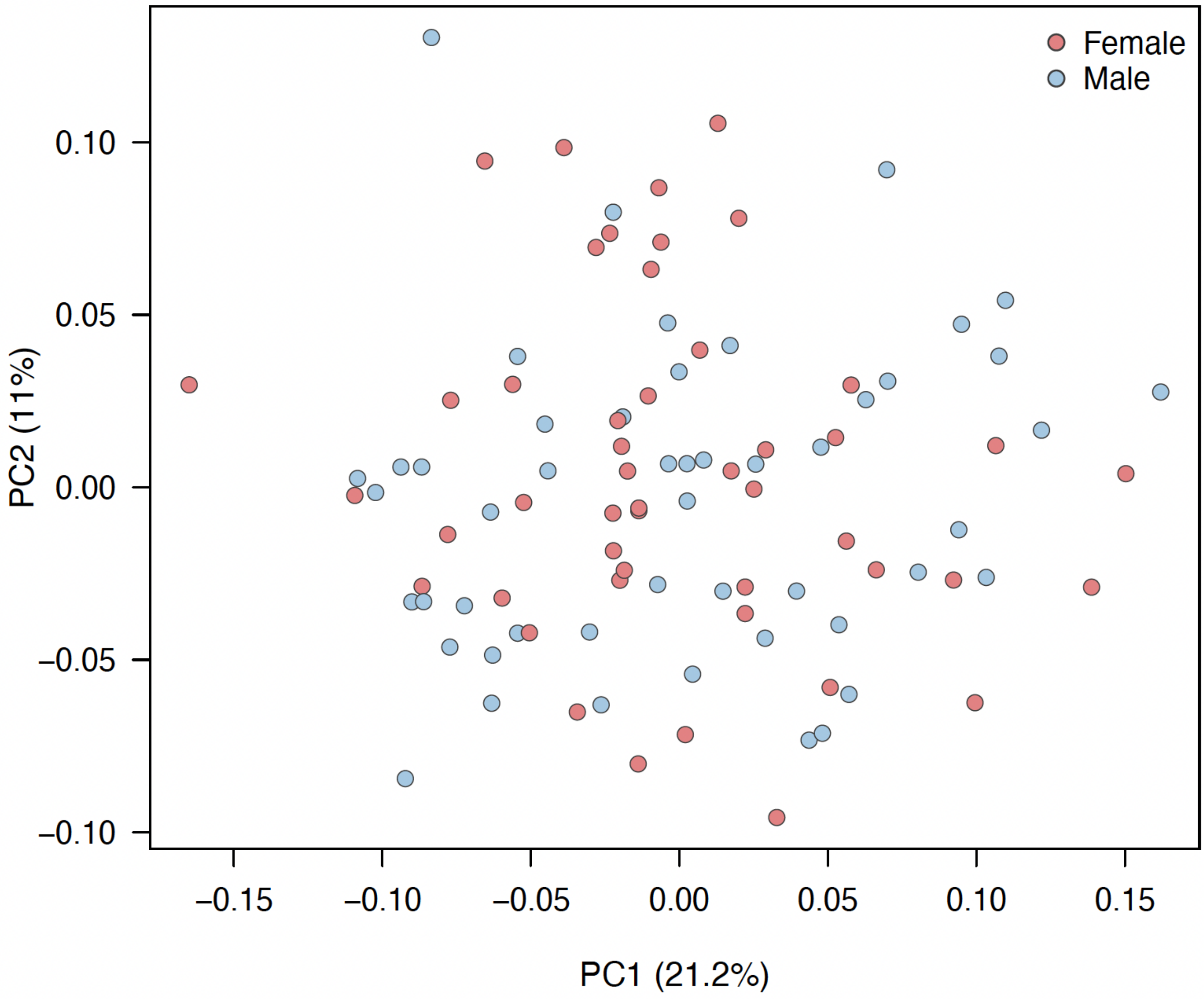


Figure S1. Principal component analysis of cochlear shape showing extensive overlap between genetically female and genetically male individuals.

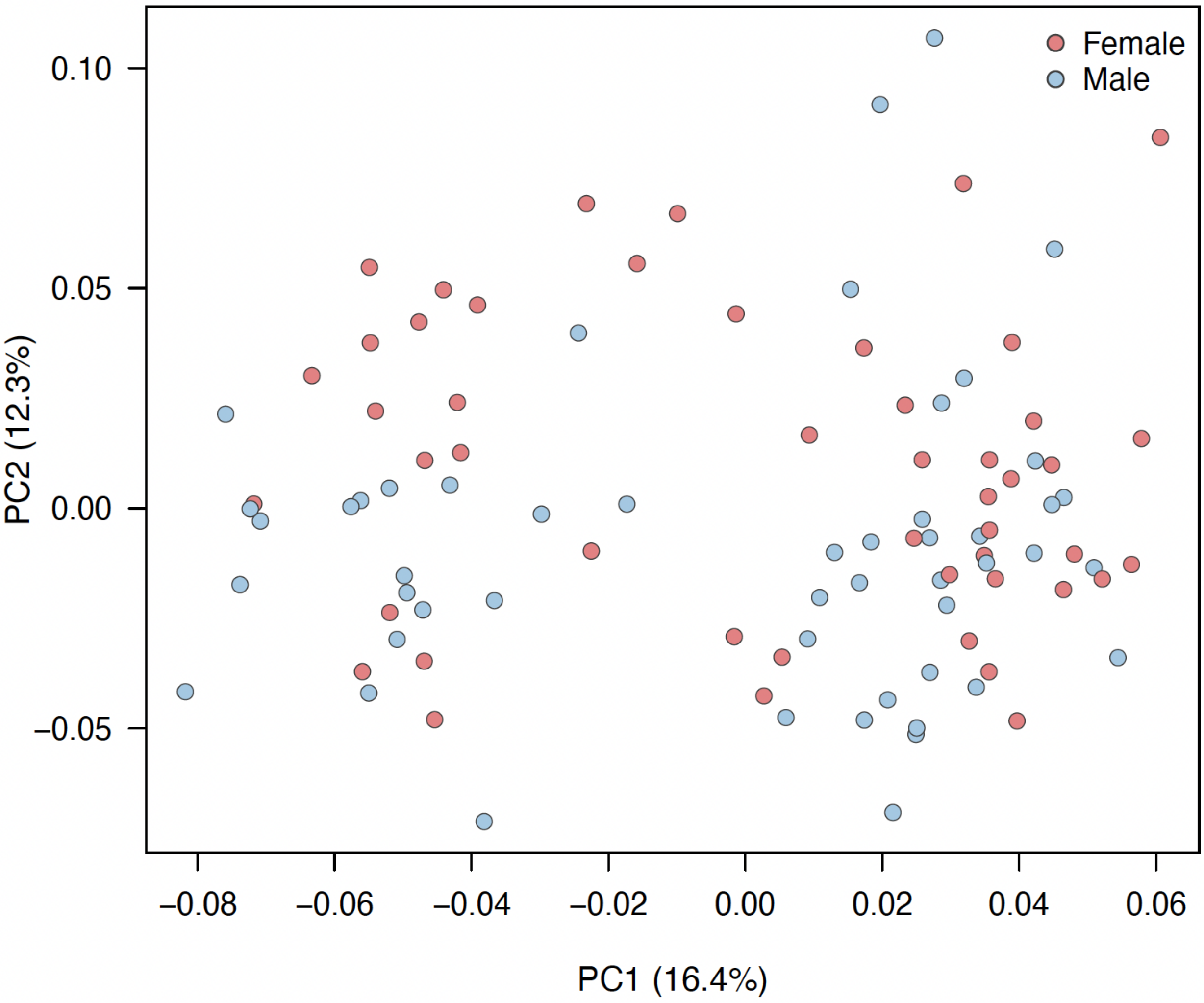


Figure S2. Principal component analysis of the combined semicircular canals showing extensive overlap between genetically female and genetically male individuals.

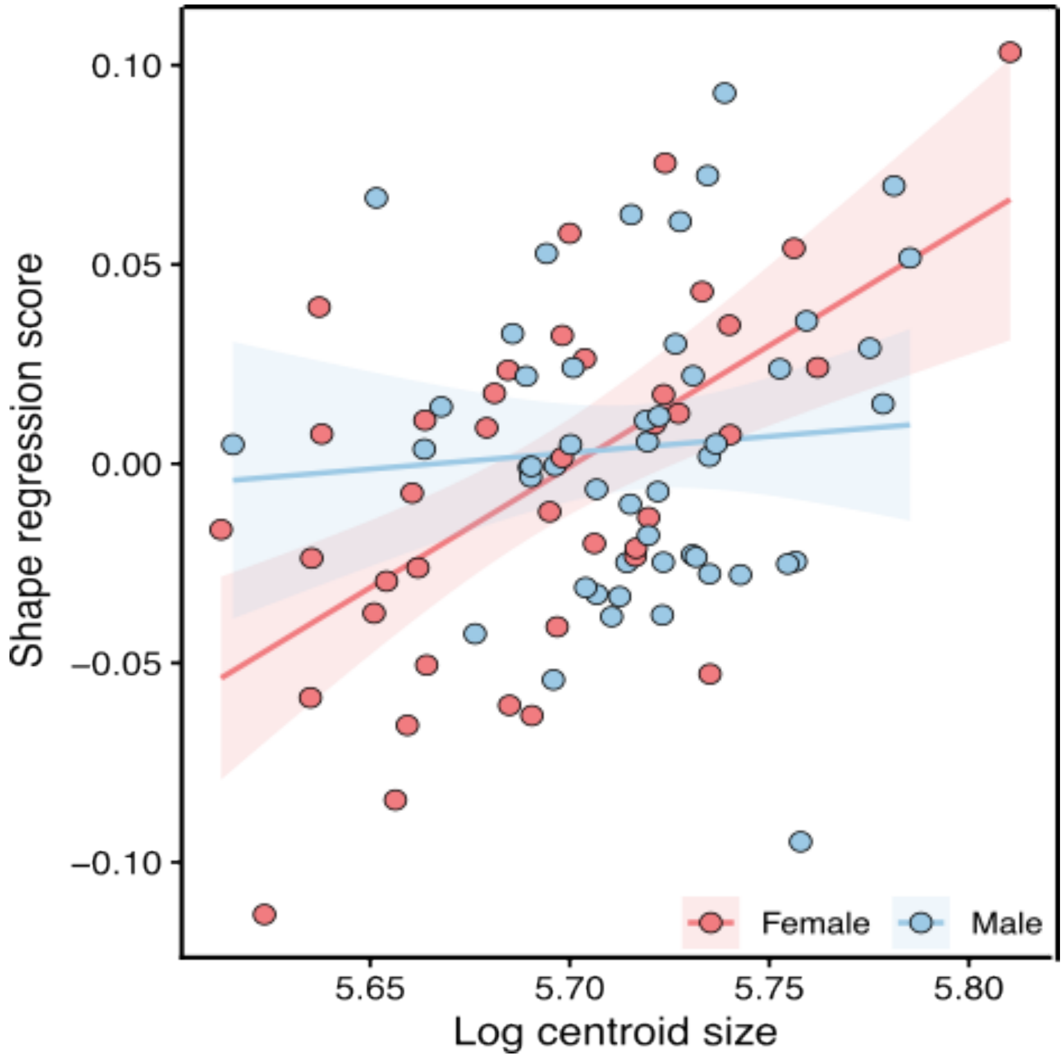


Figure S3. Allometric relationship between cochlear shape (represented by regression scores) and log-transformed centroid size. Lines represent linear regressions fitted separately for genetically female and genetically male individuals, and shaded areas indicate 95% confidence intervals. A weak allometric relationship was detected, with no evidence of sex-specific allometric trajectories.

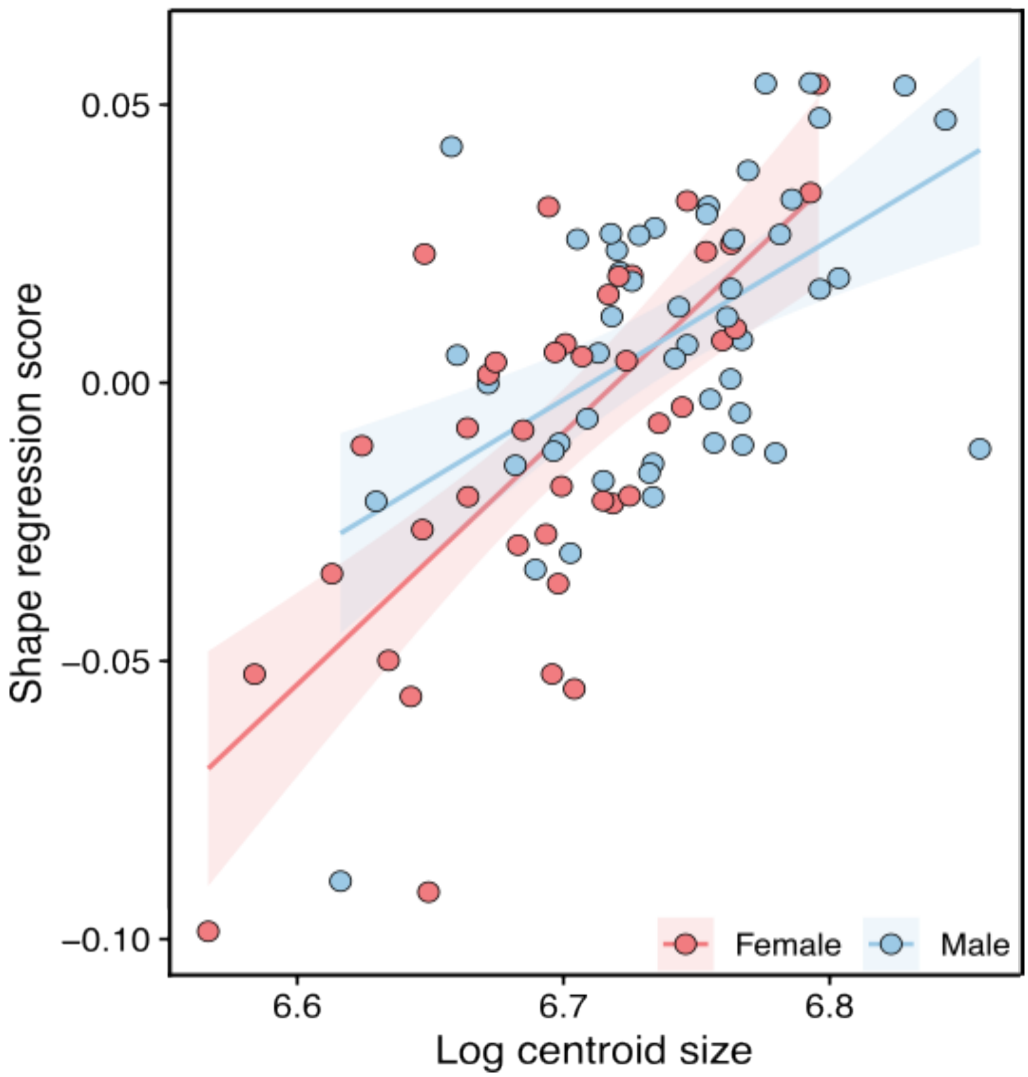


Figure S4. Allometric relationship between semicircular canal shape (represented by regression scores) and log-transformed centroid size. Lines represent linear regressions fitted separately for genetically female and genetically male individuals, and shaded areas indicate 95% confidence intervals. No significant allometric relationship or sex-specific allometric trajectories were detected.

Supplementary Table S1. Metadata for the 98 individuals included in the geometric morphometric analyses. The first column (ID) contains the unique scan identifiers used throughout image acquisition, landmarking, and geometric morphometric analyses. These identifiers are retained to facilitate reproducibility and future cross-referencing across related datasets and publications. The second and third columns (Sample and Archaeological site) correspond to the specimen designations and archaeological provenience reported by the institutions curating the collections and by the researchers who originally studied and published these materials. Laterality indicates the anatomical side (left or right) of the scanned bony labyrinth. Genomic sex refers to the genetically determined sex of each individual (F = female; M = male). Geographic region indicates the geographic provenience of each specimen, and Age group corresponds to the age category used in the present analyses. An asterisk (*) following the scan identifier indicates specimens imaged using medical computed tomography (n = 5); all other specimens were imaged using micro-computed tomography (n = 93).

| ID | Sample | Archaeological site | Laterality | Genomic Sex | Geographic Region | Age group |
|---|---|---|---|---|---|---|
| BR_708570* | Urna 8 - Crânio 51193 | Toca da Baixa dos Caboclos | right | F | Northeastern Brazil | Subadult |
| BR_708575* | Esqueleto 2 - Crânio 89098-178 | Toca do Serrote do Tenente Luiz | right | F | Northeastern Brazil | Subadult |
| MLP12_bottom | 7999 | Sta. Rosa Tastil | right | M | Northwestern Argentina | Adult |
| NOA1_bottomup | Soria 2-Rasgo17-UP13 | Mesada de Andalhuala | right | M | Northwestern Argentina | Subadult |
| MNHN_HA_10278 | 10278 | Norte Isla Tierra del Fuego | left | M | Tierra del Fuego, Chile | Adult |
| MNHN_HA_10279 | 10279 | Norte Isla Tierra del Fuego | left | M | Tierra del Fuego, Chile | Adult |
| MLP10_bottom | 1290 | Valle Chubut | right | F | Southern Patagonia, Argentina | Adult |
| MLP10_top | 7573 | Litoral norte Patagonia, Chubut | right | M | Southern Patagonia, Argentina | Adult |
| MLP7_middle | 1272 | Valle Chubut | left | F | Southern Patagonia, Argentina | Adult |
| MLP7_top | 1269 | Valle Chubut | right | M | Southern Patagonia, Argentina | Adult |
| PIMUZ_4612_left | PIMUZ_4612 | Última Esperanza | left | F | Southern Patagonia, Chile | Adult |
| CBA5c_middle | C-4-1 | Isla Barranquita 1 | left | M | Paraná Delta, Argentina | Adult |
| DE1MZA5_middle | B 651-2 | Isla Cementerio R3 | right | M | Paraná Delta, Argentina | Adult |

| | | | | | | |
|---|---|---|---|---|---|---|
| DE2_bottom | 256 | Isla Barranquita 1 | left | M | Paraná Delta, Argentina | Adult |
| DE2_middle | 236 | Isla Barranquita 1 | right | F | Paraná Delta, Argentina | Adult |
| DE2_top | Cajon428-Ind18 An-63 | Isla Barranquita 1 | left | M | Paraná Delta, Argentina | Adult |
| MLP3_bottom | 6468 | Arroyo Sarandí | right | F | Paraná Delta, Argentina | Adult |
| BA3_bottom | Ent29 m266 | Chenque 1 | left | F | Pampas, Argentina | Adult |
| MIX3_bottom | 7519 | Los Talas | left | F | Pampas, Argentina | Adult |
| MLP12_middle | 7518 | Los Talas | left | F | Pampas, Argentina | Adult |
| MLP1_top | 400 | Arroyo Chocoíi | left | M | Pampas, Argentina | Adult |
| MLP5_middle | 7497 | Los Talas | right | M | Pampas, Argentina | Adult |
| MLP8_middle | 7508 | Los Talas | left | F | Pampas, Argentina | Adult |
| MLP8_top | 7928 | Palo Blanco | left | M | Pampas, Argentina | Adult |
| PA2_middle | Ent25-m106 | Chenque 1 | right | M | Pampas, Argentina | Adult |
| PA2_top | Ent15-m15 | Chenque 1 | left | M | Pampas, Argentina | Adult |
| PA3_bottom | Ent39(2)-m37 | Chenque 1 | left | F | Pampas, Argentina | Adult |
| PA3_middle | Trq1 | La Tranquera | right | M | Pampas, Argentina | Adult |
| PA3_top | Ent6-m27 | Chenque 1 | right | M | Pampas, Argentina | Adult |
| PAMPA_AS2_6* | AS2_6 | Arroyo Seco 2 | left | F | Pampas, Argentina | Subadult |
| PAMPA_LCH_E1* | Entierro 1 | Laguna Chica | right | M | Pampas, Argentina | Adult |
| PAMPA_LCH_E4* | Entierro 4 | Laguna Chica | left | F | Pampas, Argentina | Subadult |
| CBA1_bottom | P-45 | Cerro Colorado 45 | left | F | Central Highlands, Argentina | Adult |
| CBA1_top | P-T5-I1 | Carrupachina 5 | left | F | Central Highlands, Argentina | Juvenile |
| CBA2_bottom | P-15 | Cerro Colorado 15 | left | M | Central Highlands, Argentina | Subadult |
| CBA2_middle | P-4 | Cerro Colorado 4 | left | F | Central Highlands, Argentina | Adult |
| CBA2_top | P-5 | Cerro Colorado 5 | left | F | Central Highlands, Argentina | Adult |
| CBA3_bottom | P-38 | Cerro Colorado 38 | right | F | Central Highlands, Argentina | Subadult |
| CBA3_middle | P-58 | Cerro Colorado 58 | right | F | Central Highlands, Argentina | Subadult |

| | | | | | | |
|---|---|---|---|---|---|---|
| CBA3_top | P-60 | Cerro Colorado 60 | left | M | Central Highlands, Argentina | Adult |
| CBA4 | P-7 | Cerro Colorado 7 | left | M | Central Highlands, Argentina | Adult |
| CBA5c_bottom | P-80 | Cerro Colorado 80 | left | M | Central Highlands, Argentina | Adult |
| CBA6_top | P-73 | Cerro Colorado 73 | left | F | Central Highlands, Argentina | Subadult |
| CBA7_bottom | P-LC | La Cancha | right | F | Central Highlands, Argentina | Adult |
| CBA7c_middle | P-T1-I1 | Carrupachina 1 | right | M | Central Highlands, Argentina | Adult |
| CBA7c_top | P-GT-1 | Anqas Mayo 1 | right | M | Central Highlands, Argentina | Adult |
| MIX1_bottom | P-67 | Cerro Colorado 67 | right | M | Central Highlands, Argentina | Adult |
| HIST1_bottom | M-7 | Ruinas Jesuíticas | left | F | Northern Mendoza | Adult |
| HIST1_middle | M-16 | Ruinas Jesuíticas | right | M | Northern Mendoza | Adult |
| HIST2_bottom | M-32 | La Caridad | right | F | Northern Mendoza | Adult |
| HIST2_middle | M-38 | Ruinas Jesuíticas | right | F | Northern Mendoza | Adult |
| HIST2_top | M-6 | Ruinas Jesuíticas | right | F | Northern Mendoza | Adult |
| HIST3_top | M-31 | La Merced | right | F | Northern Mendoza | Adult |
| HIST4 | M-11 | Ruinas Jesuíticas | right | M | Northern Mendoza | Adult |
| MZA10_bottomup | 5 | Barrio Ramos | right | F | Northern Mendoza | Subadult |
| MZA10_top | 1 | Las Cuevas 8 | left | M | Northern Mendoza | Subadult |
| MZA1_bottom | Las Cuevas-2 | Las Cuevas 8 | right | M | Northern Mendoza | Subadult |
| MZA1_middleup | CA19 | Capiz Alto | right | M | Northern Mendoza | Subadult |
| MZA1_top | CA8 | Capiz Alto | right | F | Northern Mendoza | Juvenile |
| MZA4_bottom | 1 | Túmulo III Uspallata | left | F | Northern Mendoza | Subadult |

| | | | | | | |
|---|---|---|---|---|---|---|
| MZA4_bottomup | 301 | Potrero Las Colonias | left | M | Northern Mendoza | Adult |
| MZA4_up | 5 | Túmulo III Uspallata | left | F | Northern Mendoza | Subadult |
| MZA4_upbottom | 3 | Túmulo III Uspallata | left | F | Northern Mendoza | Subadult |
| MZA7_bottom | 7 | Túmulo III Uspallata | left | F | Northern Mendoza | Subadult |
| MZA7_bottomup | 471c | Potrero Las Colonias | right | F | Northern Mendoza | Subadult |
| MZA7_middle | 2 | Túmulo III Uspallata | left | M | Northern Mendoza | Subadult |
| MZA7_top | 6 | Túmulo III Uspallata | left | M | Northern Mendoza | Subadult |
| MZA8_bottom | 17 | Capiz Alto | left | M | Northern Mendoza | Adult |
| MZA8_top | petroso suelto | Potrero Las Colonias | left | F | Northern Mendoza | Adult |
| MZA9_bottom | B6-22 | Barrancas B6 | right | F | Northern Mendoza | Subadult |
| MZA9_top | N5 | Natania | right | F | Northern Mendoza | Subadult |
| MIX1_middlebottom | JP1251 | Jaime Prats | right | M | Southern Mendoza | Subadult |
| MIX1_top | LaB-S5-2 | Laguna Blanca | left | F | Southern Mendoza | Adult |
| SM10_bottom | OA1-9-2 | Ojo de Agua | left | M | Southern Mendoza | Subadult |
| SM10_bottomup | OA1-4-3 | Ojo de Agua | right | F | Southern Mendoza | Subadult |
| SM11_bottom | AF1108-2 | India Muerta | right | M | Southern Mendoza | Adult |
| SM11_middle | AF513-2 | Cerro Mesa | left | M | Southern Mendoza | Adult |
| SM12_middle | AF677 | Patas de Puma | right | M | Southern Mendoza | Adult |
| SM13_middle | AF677 | Patas de Puma | right | M | Southern Mendoza | Adult |
| SM13_top | OA2-1-1-2 | Ojo de Agua | left | M | Southern Mendoza | Adult |
| SM1_bottom | Ent33 | Buta Mayil | left | M | Southern Mendoza | Adult |
| SM1_middle | Esq7 | Llancanelo | left | F | Southern Mendoza | Juvenile |

| | | | | | | |
|---|---|---|---|---|---|---|
| SM1_top | 6 | Llancanelo | right | F | Southern Mendoza | Subadult |
| SM2_bottomup | Esq15 | El Alambrado | right | F | Southern Mendoza | Adult |
| SM2_middle | Esq24-Bolsa209 | Arroyo Negro Pincheira | left | M | Southern Mendoza | Subadult |
| SM3_bottom | AF2020 | Canada Seca | right | M | Southern Mendoza | Adult |
| SM3_bottomup | AF2008 | Arroyo El Tigre | left | F | Southern Mendoza | Adult |
| SM3_top | ECH-1 | El Chacay | right | M | Southern Mendoza | Adult |
| SM3_upbottom | AF681 | Médano Puesto Diaz | right | F | Southern Mendoza | Adult |
| SM4_bottom | Puesto El Alto | Puesto El Alto | left | M | Southern Mendoza | Adult |
| SM4_up | LCAB | La Cabeza | right | M | Southern Mendoza | Adult |
| SM4_upbottom | AF2074 | Pto. La Huertita | left | F | Southern Mendoza | Subadult |
| SM5_upbottom | AF505 | La Matancilla | left | M | Southern Mendoza | Adult |
| SM6_bottom | AF2032 | El Nihuil | left | M | Southern Mendoza | Adult |
| SM6_bottomtop | AF2015 | Médano Puesto Diaz | left | M | Southern Mendoza | Adult |
| SM6_top | AF500 | Rincol del Atuel | right | M | Southern Mendoza | Adult |
| SM7_bottom | AF1103 | Campo Las Julias - Las Toscas | right | M | Southern Mendoza | Adult |
| SM9_top | JP1178-Cr36 | Jaime Prats | right | M | Southern Mendoza | Juvenile |

Supplementary Table S2. Archaeological sites represented in the study and their geographic provenience. References correspond to the primary archaeological publications describing each site, assemblage, or associated human remains.

| Archaeological site | Geographic region | Archaeological reference(s) |
|---|---|---|
| Toca da Baixa dos Caboclos | Northeastern Brazil | Guidon et al. (1998); Mendonça de Souza et al. (2002) |
| Toca do Serrote do Tenente Luiz | Northeastern Brazil | Cunha (2014) |
| Sta Rosa Tastil | Northwestern Argentina | Cigliano (1973) |
| Mesada de Andalhuala | Northwestern Argentina | Spano et al.(2014) |
| Norte Isla Tierra del Fuego | Tierra del Fuego, Chile | Galland & Friess (2016) |
| Valle Chubut | Southern Patagonia, Argentina | Lehmann-Nitsche (1910) |
| Litoral norte Patagónico | Southern Patagonia, Argentina | Lehmann-Nitsche (1910) |
| Última Esperanza | Southern Patagonia, Chile | Menéndez et al. (2025); Schmelzle et al. (2025) |
| Isla Barranquita 1 | Paraná Delta, Argentina | Cocco et al. (2004) |
| Isla Cementerio R3 | Paraná Delta, Argentina | Cocco et al. (2004) |
| Arroyo Sarandí | Paraná Delta, Argentina | Ramos Van Rap et al. (2016) |
| Chenque 1 | Pampas, Argentina | Berón (2018) |
| Los Talas | Pampas, Argentina | Del Papa et al. (2020) |
| Arroyo Chocorí | Pampas, Argentina | Politis & Bonomo (2011) |
| Palo Blanco | Pampas, Argentina | Del Papa et al. (2020) |
| La Tranquera | Pampas, Argentina | Berón et al. (2015) |
| Arroyo Seco 2 | Pampas, Argentina | Politis et al. (2014) |
| Laguna Chica | Pampas, Argentina | Scheifler et al. (2024, 2025) |
| Cerro Colorado 45 | Central Highlands, Argentina | Diaz & Recalde (2025); Recalde et al. (2024); Tissera et al. (2019) |
| Carrupachina 5 | Central Highlands, Argentina | Rivero et al. (2015) |
| Cerro Colorado 15 | Central Highlands, Argentina | Diaz & Recalde (2025); Recalde et al. (2024); Tissera et al. (2019) |

| | | |
|---|---|---|
| Cerro Colorado 4 | Central Highlands, Argentina | Diaz & Recalde (2025); Recalde et al. (2024); Tissera et al. (2019) |
| Cerro Colorado 5 | Central Highlands, Argentina | Diaz & Recalde (2025); Recalde et al. (2024); Tissera et al. (2019) |
| Cerro Colorado 38 | Central Highlands, Argentina | Diaz & Recalde (2025); Recalde et al. (2024); Tissera et al. (2019) |
| Cerro Colorado 58 | Central Highlands, Argentina | Diaz & Recalde (2025); Recalde et al. (2024); Tissera et al. (2019) |
| Cerro Colorado 60 | Central Highlands, Argentina | Diaz & Recalde (2025); Recalde et al. (2024); Tissera et al. (2019) |
| Cerro Colorado 7 | Central Highlands, Argentina | Diaz & Recalde (2025); Recalde et al. (2024); Tissera et al. (2019) |
| Cerro Colorado 80 | Central Highlands, Argentina | Diaz & Recalde (2025); Recalde et al. (2024); Tissera et al. (2019) |
| Cerro Colorado 73 | Central Highlands, Argentina | Diaz & Recalde (2025); Recalde et al. (2024); Tissera et al. (2019) |
| La Cancha | Central Highlands, Argentina | Tissera (2014) |
| Carrupachina 1 | Central Highlands, Argentina | Rivero et al. (2015) |
| Anqas Mayo 1 | Central Highlands, Argentina | Tissera (2024) |
| Cerro Colorado 67 | Central Highlands, Argentina | Diaz & Recalde (2025); Recalde et al. (2024); Tissera et al. (2019) |
| Ruinas Jesuíticas | Northern Mendoza | Chiavazza et al. (2015) |
| La Caridad | Northern Mendoza | Chiavazza et al. (2015) |
| La Merced | Northern Mendoza | Chiavazza et al. (2015) |
| Barrio Ramos | Northern Mendoza | Duran et al. (2018) |
| Las Cuevas 8 | Northern Mendoza | Unpublished |
| Capiz Alto | Northern Mendoza | Duran & Novellino (1999-2000) |
| Túmulo III Uspallata | Northern Mendoza | Rusconi (1962); Barberena et al. (2026) |
| Potrero Las Colonias | Northern Mendoza | Guevara et al. (2022) |
| Barrancas B6 | Northern Mendoza | Novellino et al. (2013); Barberena et al. (2017) |

| | | |
|---|---|---|
| Natania | Northern Mendoza | Unpublished |
| Jaime Prats | Southern Mendoza | Gil et al. (2020); Peralta et al. (2024) |
| Laguna Blanca | Southern Mendoza | Gil et al. (2020); Peralta et al. (2024) |
| Ojo de Agua | Southern Mendoza | Gil et al. (2020); Peralta et al. (2024) |
| India Muerta | Southern Mendoza | Gil et al. (2020); Peralta et al. (2024) |
| Cerro Mesa | Southern Mendoza | Gil et al. (2020); Peralta et al. (2024) |
| Patas de Puma | Southern Mendoza | Gil et al. (2020); Peralta et al. (2024) |
| Buta Mayil | Southern Mendoza | Gil et al. (2020); Peralta et al. (2024) |
| Llancanelo | Southern Mendoza | Gil et al. (2020); Peralta et al. (2024) |
| El Alambrado | Southern Mendoza | Gil et al. (2020); Peralta et al. (2024) |
| Arroyo Negro Pincheira | Southern Mendoza | Gil et al. (2020); Peralta et al. (2024) |
| Cañada Seca | Southern Mendoza | Gil et al. (2020); Peralta et al. (2024) |
| Arroyo El Tigre | Southern Mendoza | Gil et al. (2020); Peralta et al. (2024) |
| El Chacay | Southern Mendoza | Gil et al. (2020); Peralta et al. (2024) |
| Médano Puesto Díaz | Southern Mendoza | Gil et al. (2020); Peralta et al. (2024) |
| Puesto El Alto | Southern Mendoza | Gil et al. (2020); Peralta et al. (2024) |
| La Cabeza | Southern Mendoza | Gil et al. (2020); Peralta et al. (2024) |
| Puesto La Huertita | Southern Mendoza | Gil et al. (2020); Peralta et al. (2024) |

| | | |
|---|---|---|
| La Matancilla | Southern Mendoza | Gil et al. (2020); Peralta et al. (2024) |
| El Nihuil | Southern Mendoza | Gil et al. (2020); Peralta et al. (2024) |
| Rincón del Atuel | Southern Mendoza | Gil et al. (2020); Peralta et al. (2024) |
| Campo Las Julias – Las Toscas | Southern Mendoza | Gil et al. (2020); Peralta et al. (2024) |

Supplementary Table S3. Anatomical definition of the 64 landmarks used to characterize the shape of the human bony labyrinth.

| |
|---|
| **Cochlea** |
| 1. Cochlear center |
| 2. Apex of the cochlea |
| 3. Distal point of the apical turn of the cochlea |
| 4. Inner point of the middle turn of the cochlea |
| 5. Proximal point of the middle turn of the cochlea |
| 6. Outer point of the middle turn of the cochlea |
| 7. Distal point of the middle turn of the cochlea |
| 8. Inner point of the basal turn of the cochlea |
| 9. Proximal point of the basal turn of the cochlea |
| 10. Outer point of the basal turn of the cochlea |
| 11. Distal point of the basal turn of the cochlea |
| 13. Cochlear center |
| 14. Proximal point of the lower portion of the middle turn of the cochlea |
| 15. Outer point of the middle turn of the cochlea |
| 16. Distal point of the upper portion of the basal turn of the cochlea |
| 17. Inner point of the basal turn of the cochlea |
| 18. Proximal point of the lower portion of the basal turn of the cochlea |
| **Vestibular system** |
| 12. Saccular apex |
| 19. Basal, lowest point of the LSCC |

| |
|---|
| 20. Left internal point of the LSCC |
| 21. Upper internal point of the LSCC |
| 22. Right internal point of the LSCC |
| 23. Left external point of the LSCC |
| 24. Highest point of the LSCC |
| 25. Right external point of the LSCC |
| 26. Deeper point of the LSCC |
| 27. Left internal point of the LSCC |
| 28. Upper internal point of the LSCC |
| 29. Most prominent point of the LSCC protuberance |
| 30. Left external point of the LSCC |
| 31. Highest external point of the LSCC |
| 32. Right external point of the LSCC |
| 33. Lowest point of the vestibular wall |
| 34. Left internal point of the ASCC |
| 35. Upper internal point of the ASCC |
| 36. Right internal point of the ASCC |
| 37. Left external point of the ASCC |
| 38. Highest external point of the ASCC |
| 39. Right external point of the ASCC |
| 40. Deeper point of the ASCC |
| 41. Left internal point of the ASCC |
| 42. Upper internal point of the ASCC |
| 43. Most prominent point of the ASCC protuberance |
| 44. Left external point of the ASCC |
| 45. Highest external point of the ASCC |
| 46. Right external point of the ASCC |
| 47. Meeting point of the vestibular wall and the edges of the common cross, and the base of the lateral canal. |
| 48. Left internal point of the PSCC |
| 49. Upper internal point of the PSCC |

| |
|---|
| 50. Right internal point of the PSCC |
| 51. Left external point of the PSCC |
| 52. Highest point of the PSCC |
| 53. Right external point of the PSCC |
| 54. Most prominent internal point of the PSCC protuberance |
| 55. Left internal point of the PSCC |
| 56. Upper internal point of the PSCC |
| 57. Right internal point of the PSCC |
| 58. Most prominent external point of the PSCC* |
| 59. Left external point of the PSCC |
| 60. Highest external point of the PSCC |
| 61. Right external point of the PSCC |
| 62. Meeting point of the end of the basal turn of the cochlea and the PSCC |
| 63. Meeting point between the ASCC and the PSCC on the edge of the common cross. |
| 64. Meeting point between the ASCC and the LSCC |
